# Clostridium ablates tumors by degrading collagen mesh and inducing ferroptosis

**DOI:** 10.1101/2024.05.15.594430

**Authors:** Fengmin Yang, Lu Meng, Huan Chen, Xinyue Wang, Mengmeng Zhang, Zhenping Cao, Yan Pang, Jinyao Liu

## Abstract

The supportive role of extracellular matrix in tumor growth and the drug resistance of tumor cells stand the major challenges in treating cancer. Here, we report the discovery of an oncolytic Clostridium strain, named RJ-1, which simultaneously expresses collagenases and produces iron-containing particles through intrabacterial mineralization. The secreted collagenases degrade pre-existing collagen mesh and show a collagen concentration-dependent degradation feature. Iron-containing particles released from RJ-1 by autolysis generate ferrous ions in the lysosomal acid environment following cell uptake and subsequently induce ferroptosis via the Fenton reaction. Due to selective germination in tumor sites, systemic administration of RJ-1 spores synchronously destructs dense tumor extracellular matrix and causes non-apoptotic tumor cell death. In collagen-dense tumor-bearing mice, a single intravenous injection achieves a favorable tolerance and rapid oncolysis of giant tumors with a size of ∼1000 mm^3^. This work finds a dual collagenase-expressing and iron-concentrating bacterium, proposing an alternative strategy to combat tumors.

## Main

Both the supportive role of extracellular matrix (ECM) in tumor growth and the drug resistance of tumor cells constitute the major challenges in cancer treatment ^1,2^. As a primary component of extracellular matrix and a unique nutritional source, collagen not only facilitates the proliferation and invasion of tumor cells through signal transduction, but also inhibits the infiltration and activation of immune cells ^1^. In addition, inherent apoptotic resistance as well as resistance acquired by the accumulation of mutations lead to considerable tolerance of tumor cells toward drug-induced cell death pathways ^2,3^. Given these facts, strategies capable of controlling the deposition of collagen fibers ^4^ and inducing non-apoptotic cell death are highly desirable for developing effective tumor therapies ^5^.

Utilizing either small-molecule agents to suppress collagen generation and disturb its cross- linking process ^4^ or collagenases to degrade and destruct collagen fibers represents a promising strategy to elicit antitumor responses ^6^. While, the use of small-molecular inhibitors and collagenases as well as their nanoparticular formulations can result in non-specific degradation of systemic collagen, posing a potential risk of hemorrhaging in health organs ^7,8^. As an independent cell death pathway, ferroptosis has attracted substantial attention to addressing tumor resistance against drug-induced apoptosis ^3^. Yet, the process of ferroptosis is highly context-dependent ^9^, limiting the ability to target specific proteins in ferroptosis pathway across different types of cancer^5^. Meanwhile, despite the employment of iron-containing nanoparticles is able to directly increase intracellular ferric ion concentration and corresponding activity of the Fenton reaction ^10^, this non-specific pattern inevitably induces undesired side effects due to uncontrollable distribution of nanoparticles in vivo ^11,12^.

In this study, we find an oncolytic *Clostridium bifermentans* strain, termed RJ-1, which can simultaneously express collagenases and synthesize iron-containing particles (INPs) via intrabacterial mineralization. Due to its living characteristic, RJ-1 degrades pre-existing collagen mesh and displays a collagen concentration-dependent degradation manner. RJ-1 also releases INPs through autolysis, thereby inducing Fenton reaction-mediated cell ferroptosis given the uptake of these INPs generates ferrous ions in the lysosomal acid environment. Benefiting from selective germination in the hypoxic and immunosuppressive tumor sites, systemic administration of RJ-1 spores synchronously depletes intratumoral ECM and elicits non-apoptotic tumor cell death. In an orthotopic triple-negative breast carcinoma (TNBC) murine model with collagen-dense ECM, a single intravenous dose achieves a satisfactory tolerance and immediate oncolysis of tumors even with a size up to ∼1000 mm^3^. The ability of RJ-1 to lyse tumors is further supported by using an orthotopic pancreatic ductal adenocarcinoma (PDAC) mouse model. Our work discloses a dual collagenase-expressing and iron-concentrating Clostridium strain, providing an effective yet safe platform to combat cancer by destructing tumor stroma and circumventing drug resistance.

### Isolation and characterization of RJ-1

By using skim milk plates, we successfully isolated a collagenase-producing Clostridium strain, which was identified as *Clostridium bifermentans* through 16S ribosomal RNA sequencing and named RJ-1 (Fig. S1). After incubation on differential clostridial agar plates under an anaerobic condition, RJ-1 colonies turned black, which was associated with ferrous sulfide (FeS) production (Fig. S2A). RJ-1 exhibited β-hemolysis on Columbia blood agar plates (Fig. S2B), indicating its capability to acquire iron sources from erythrocytes in vivo ^13,14^. Interestingly, branch-like colonies were formed on brain heart infusion (BHI) plates supplemented with 1% agar and accompanied with further extension of the colonies and the appearance of umbilical fossa-like hollow area in the center with incubation time elongating to 10 days (Fig. S2C). This phenomenon suggested the swarming motility and autolytic activity of the isolated RJ-1 ^15^. As indicated in transmission electron microscopy (TEM) images shown in Fig. S2D, RJ-1 existed in two forms including rod-shaped vegetative cells and oval spores, which could be separated by using density gradient centrifugation and readily distinguished under a dark field microscopy (Fig. S2E).

### Degradation of collagen in vitro by collagenase secreted from RJ-1

Compared to typical *Enterococcus faecalis*, *Bacillus cereus*, *Escherichia coli* Nissle 1917, and *Salmonella typhimurium* VNP20009, RJ-1 shaped the largest diameter of the transparent zone on skim milk plates (Fig. S2F), indicating its highest level of collagenase production ^16^. As depicted in Fig. 1A, RJ-1 almost completely degraded the acid-soluble collagen hydrogel and left a liquefied cavity. To assess the impact of RJ-1 on ECM, fibroblast-secreted ECM components were collected and co-cultured with RJ-1 ^17^. Following 12 hours incubation, there was a significant elimination of collagen, while the collagen fibers treated with phosphate buffer saline (PBS) retained intact (Fig. 1B, C). Furthermore, revealed by both sodium dodecyl sulfate polyacrylamide gel electrophoresis (SDS-PAGE) and gelatin zymography assay (Fig. 1D, E), RJ-1 secreted a serial of collagenases with molecular weights ranging from 35 to 100 kilodalton (kDa). These protein-level results were supported by the presence of collagenase-associated proteases through RJ-1 genome sequencing (Table S1).

**Fig 1.**
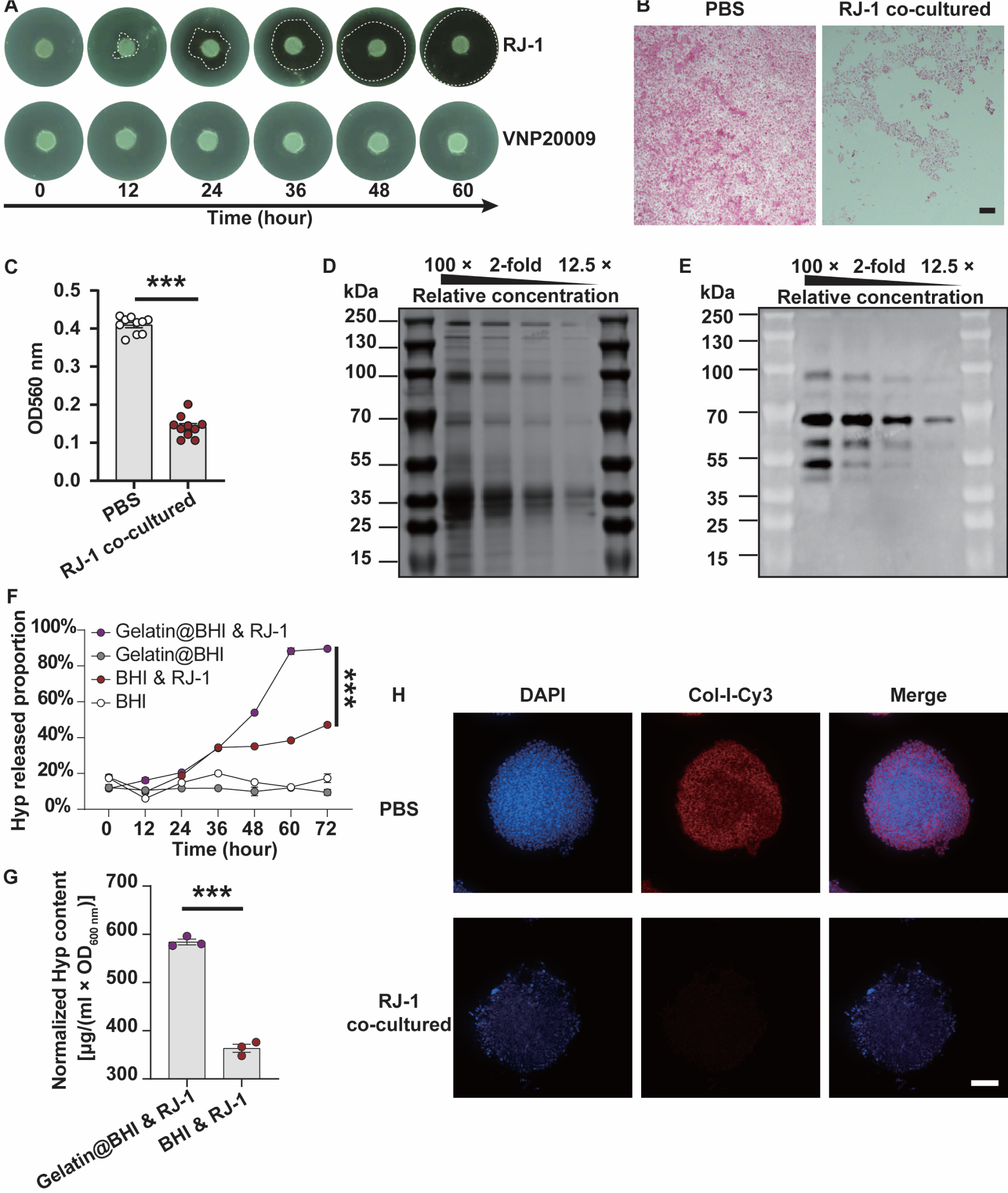
Degradation of collagen in vitro by collagenase secreted from RJ-1. (A) Digital photos of acid soluble collagen hydrogel after incubation with RJ-1 for the indicated time points. White dotted circles outline the cavities formed by degradation. (B) Confocal images of fibroblast-secreted collagen after incubation with PBS and RJ-1 for 12 hours, respectively. Collagen was stained with picrosirius red. Scale bar: 100 μm. (C) Quantification of the remained collagen content in the extracellular matrix of fibroblasts. The absorbance was measured at OD_560 nm_ to determine the extent of degradation (*n* = 10). (D) SDS-PAGE and (E) gelatin zymography assay of the secreted proteins in the culture supernatant of RJ-1. Two-fold serial dilution was added from left to right. (F) Released of Hyp by RJ-1 in BHI or BHI containing gelatin (*n* = 3). (G) Released content of Hyp normalized to OD_600 nm_ (*n* = 3). (H) Col-I immunofluorescence of cell spheroids after co-incubation with PBS or RJ-1 for 16 hours (*n* = 3). Scale bar: 100 μm. Cell nuclei were stained with 2-(4-amidinophenyl)-6-indolecarbamidine dihydrochloride (DAPI). For statistical analysis of picrosirius red absorbance and normalized Hyp content, students’ *t* test was performed for *p* value generation. For statistical analysis of Hyp released proportion, one-way ANOVA was used and Tukey test was performed for *p* value generation. ***, *p* < 0.001. Error bar refers to mean value ± standard error of mean (SEM).

Collagenase activity assay was performed in the presence or absence of gelatin, a processed derivative of collagen, and the total collagenase activity was quantified basing on hydroxyproline (Hyp) content, a characteristic product of collagen degradation ^18^. Expectedly, when RJ-1 was provided with gelatin, the proportion of Hyp release increased rapidly after 36 hours incubation and reached up to ∼90% at 72 hours, which was 1.9-times higher than the gelatin-free control (Fig. 1F). We examined the collagenase production by measuring the content of released Hyp and found that the addition of gelatin increased per-cell collagenase production by 1.46-times (Fig. 1G). The variation in collagenase expression at the single-cell level might arise from the sensing of external substrate concentrations and its coupling to collagenase expression regulation ^19^. This was distinguished from genetically engineered heterogeneous protein expression and could aid in sufficient destruction of the ECM within tumors possessing high abundance of collagen fibers ^20^. To explore the activity of RJ-1 in the tumor microenvironment, we established an in vitro three-dimensional (3D) culture to mimic dense tumors. Considering the lack of hypoxic region and necrotic core in tumor spheroids to support sufficient growth of anaerobic bacteria ^21–23^, co-culture of RJ-1 with tumor spheroids composed of mouse 4T1 mammary carcinoma cells and NIH/3T3 fibroblasts was conducted under a hypoxic atmosphere and bacterial broth-supplemented condition ^24^. This protocol was adapted from a previously established method for co-culture of anaerobic bacteria and epithelial cells ^25^. As expected, the collagen fibers in these tumor spheroids were completely disrupted by RJ-1, as indicated by dramatically reduced Cy3 signal of type I collagen (Col-I) and a rougher morphology after co-culture (Fig. 1H), suggesting the involvement of collagenases ^26^. Additionally, the fluorescent intensity of nuclei became remarkably weak, likely due to nucleic acid degradation mediated by bacterial DNase (Table S2).

Next, we attempted to visualize whether this collagen degradation resulted in an enlargement of collagen mesh pore size. The penetration depths of small-molecular Hoechst 33342 and 1,1’- dioctadecyl-3,3,3’,3’-tetramethylindocarbocyanine perchlorate-labeled liposomes in the tumor spheroids were examined by confocal imaging. Despite the weakened signal of cell nuclei caused by RJ-1 co-culture, the blue fluorescence of Hoechst 33342 was evenly distributed throughout the entire tumor spheroid, even the central core. On the contrary, without RJ-1, the fluorescence signals were mostly confined in the periphery (Fig. S3A). Similar results of penetration were observed by using liposomes. As evidenced by the bright red fluorescence in closer proximity to central regions, RJ-1 deepened the penetration of liposomes in the tumor spheroids (Fig. S3B). These results demonstrated that after co-culture with RJ-1, the collagen network within the tumor spheroids formed larger pores, which were sufficient even for the penetration of large-sized liposomes.

The ability of RJ-1 to degrade collagen was further characterized by simulating the remote effect of bacteria colonized in the necrotic area on the non-necrotic zone ^27^. RJ-1 was embedded within BHI agar, which was attached to the bottom of the plate to create a nutrient concentration gradient against the tumor spheroids. Within 24 hours, although the VNP20009 control grew significantly in agar and induced turbidity in the field of view, the shape of the tumor spheroids remained nearly unchanged. In contrast, the proliferation of RJ-1 led to rapid disintegration of the tumor spheroids, suggesting a strong collagenase activity and the potential to ablate tumors by the remote effect on viable tumor tissue (Fig. S3C). Therefore, we speculated that the in vivo antitumor effect of RJ-1 might differ from tumor growth delayed by VNP20009 ^28^, but induce a rapid oncolytic phenomenon of an existing tumor.

### RJ-1 mediated in vivo degradation of intratumoral collagen

To investigate the in-situ collagenase production and collagen degradation in tumor sites, RJ-1 was intratumorally injected using a TNBC mouse model. Taking into account the non-specific response against bacterial components or activities, such as different immune reactivities to Gram-positive and Gram-negative bacteria ^29^, both VNP20009 and *Bacillus cereus* were used as controls to evaluate collagen degradation. To ensure similar vitality and active metabolism, all of the bacteria used for these experiments were collected at the logarithmic growth phase. We first checked the survival of anaerobic RJ-1 in the tumor site 10 days post-injection. As shown in Fig. S4A, numerous RJ-1 with partial aggregation were clearly observed in the periphery of tumor necrotic region, indicating the preservation of vitality and swarming behavior ^30^. As evidenced by histochemistry results along with quantification analysis of picrosirius red staining of collagen fibers, RJ-1 significantly reduced collagen content around its colonization site within the tumor (Fig. S4B-C). As expected, *Bacillus cereus,* with a weaker collagenase activity, did not show a comparable collagen degradation effect. Surprisingly, we found a significant increase of local collagen content induced by VNP20009 in both necrotic and viable areas. This could be ascribed to the fibrosis that occurs when the immune system responds against bacterial components and may have a negative impact on the treatment of tumors with dense ECM ^31^.

We then investigated the performance of RJ-1 in TNBC mice by systemic administration. Given the favorable safety and preferential germination in tumor sites ^32^, the tumor-bearing mice were intravenously injected with RJ-1 spores and tumor tissues were harvested 10 days post-injection. As indicated by picrosirius red staining in Fig. 2A, RJ-1 exhibited the capability to degrade tumor collagen in a dose-dependent manner. Especially, when 10^8^ RJ-1 spores were injected, only a few sparsely arranged and disordered fibers were observed. As Col-I is the predominant collagen type in breast cancer and an important ECM component that promotes tumor development ^33–35^, we examined whether the decreased density in picrosirius red staining was accompanied by the reduction of Col-I. Indeed, western blot analysis demonstrated that Col-I was barely visible in the high-dose group (Fig. 2B). Moreover, quantitative analysis of the total collagen through Hyp measurement showed that there was a reduction to 65% of the original content when mice were treated with 10^8^ RJ-1 spores (Fig. 2C), comparable to the short-term effect achieved by a single dose of intratumoral collagenase injection ^36^. Different from collagenases that suffer rapid inactivation in a physiological setting ^37^, systemic administration of RJ-1 spores could provide a prolonged effect on intratumoral collagen reduction.

**Fig 2.**
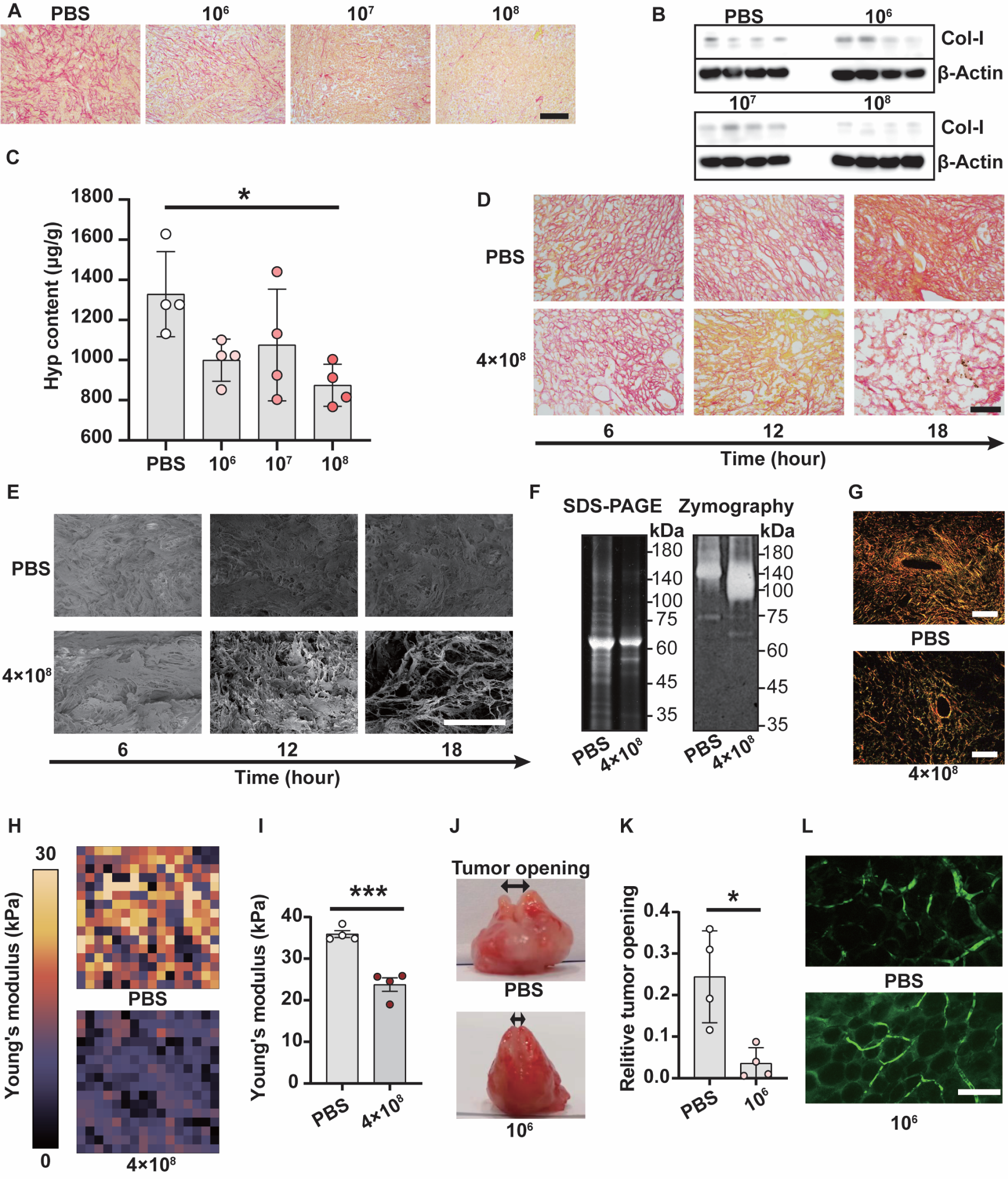
RJ-1 mediated in vivo degradation of intratumoral collagen. (A) Picrosirius red staining of tumor tissues sectioned 10 days after treating mice with RJ-1 spores intravenously at different doses or PBS. Scale bar: 100 μm. (B) Col-I content in tumor tissues sampled 10 days after treating mice with RJ-1 spores intravenously at different doses or PBS. β-Actin was used as a control. (C) Total amount of tumor collagen assessed by Hyp content 10 days after intravenous injection of different doses of RJ-1 spores or PBS (*n* = 4). (D) Picrosirius red staining of tumor tissues sectioned at the indicated time points after intravenous injection with 4 × 10^8^ RJ-1 spores or PBS. Scale bar: 100 μm. (E) SEM images of the collagen mesh structure of tumor tissues collected at the indicated time points after intravenous injection with 4 × 10^8^ RJ-1 spores or PBS. Scale bar: 50 μm. (F) Coomassie brilliant blue staining (left) and gelatin zymography analysis (right) of equivalent protein content from tumor tissues sampled 18 hours after injection with of 4 × 10^8^ RJ-1 spores or PBS. (G) Polarized light microscopy images of picrosirius red stained tumor tissues sampled 18 hours post treatment with 4 × 10^8^ RJ-1 spores or PBS. Scale bar: 200 μm. (H) Young’s modulus of tumor tissues detected by using the force spectroscopy mode of atomic force microscope. (I) Statistical analysis of the Young’s modulus of tumor tissues (*n* = 4). (J) Digital photos of the opening experiment of entire tumors 10 days after intravenous injection with 10^6^ RJ-1 spores or PBS. (K) Quantification of the relative tumor opening (*n* = 4). (L) Fluorescence imaging of dextran-FITC spreading from blood vessels to the tumor at day 10 post injection with 10^6^ RJ-1 spores or PBS. Images were captured by in vivo imaging system. Scale bar: 100 μm. For statistical analysis of Hyp content, ordinary one-way ANOVA was used and Tukey test was performed for multiple comparison and *p* value generation. For tumor opening experiment and Young’s modulus analysis, unpaired student’s *t* test was used to generate *p* value. *, *p* < 0.05; ***, *p* < 0.001. Error bar refers to mean value ± SEM.

As TNBC grows, the total amount of collagen fibers accumulates continuously, with their proportion gradually increasing ^36^. During this process, newly synthesized collagens integrate into the mature collagen mesh, undergoing modifications and cross-linking, thus exhibiting a higher stiffness ^38^, a less collagenase susceptibility ^39^, and a predominant contribution to tumor elasticity ^36,40^. We hence investigated whether RJ-1 could destruct pre-existing collagen mesh. Following a single intravenous injection of 4 × 10^8^ RJ-1 spores, tumor samples were collected at different time points for further analysis. As indicated by picrosirius red staining in Fig. 2D, the degradation of collagen fibers in the tumor started at 12 hours post-injection and apparent fractures and breakages were observed with time increasing to 18 hours. In addition, TEM images presented that these destructions were co-localized with RJ-1 vegetative cells (indicated by red arrows in Fig. S5A) and formed significant contrast with tightly arranged collagen fibers in the non-necrotic area (indicated by yellow arrows in Fig. S5B). Consistently, 3D analysis by scanning electron microscopy (SEM) displayed dynamic changes in the substantial structure of isolated collagen meshes (Fig. 2E, Fig. S5C). Furthermore, widespread disappearance of large-molecular weight proteins was found in the tumor, which could be attributed to the proteolytic activities of RJ-1 (Fig. 2F). To verify the rapid degradation of Col-I in the tumor after RJ-1 treatment, polarized light imaging was performed. As shown in Fig. 2G, in good agreement with previous results, the tightness of the arranged Col-I fibers was reduced significantly. These findings suggested an efficient disruption ability of RJ-1 against pre-existing collagen mesh in TNBC.

Collagen deposition also increases interstitial fluid pressure, leading to poor blood perfusion into tumors. Thus, we studied the mechanical property of tumors after treatment with RJ-1. Using the force spectroscopy mode of atomic force microscope, we carried out elastic modulus analysis of tumor sections and found that RJ-1 treatment rapidly led to a notable reduction in Young’s modulus, particularly at the highest injection dose of 4 × 10^8^ spores (Fig. 2H, I). Even though there was no significant reduction in the total collagen content in the lowest injection dose (10^6^) group, the interstitial fluid pressure of the tumor was significantly lowered (Fig. 2J, K). Meanwhile, the transvascular spreading of Dextran-FITC was strongly enhanced, evidenced by the accumulation of fluorescent signal in the extravascular area of the tumor (Fig. 2L), indicating the presence of larger pores within tumor ECM ^41,42^.

We further investigated whether RJ-1 could trigger collagen degradation in a PDAC mouse model. As shown in Fig. S6a, following a single dose of RJ-1 spores, the decline of collagen in the tumor became evident from 48 hours onwards. To validate the correlation between spore germination and collagen degradation, we conducted immunofluorescence staining of collagen fibers and in-situ hybridization staining of vegetative cells simultaneously. As illustrated in Fig. S7-S8, extensive collagen degradation happened as soon as the spores were injected, resulting in negligible Col-I residue at 14 days post-injection. Basing on fluorescence in-situ hybridization, substantial vegetative cells with a rod-shaped morphology were observed within the tumor on day 2, accompanied by the reduction of Col-I in the identical area. Interestingly, the fluorescence signal diminished on day 7 but reappeared in limited regions with a cell-like shape on day 14. As similarly observed in TBNC (Fig. S9), this was likely ascribed to re-entering into the sporulation process of RJ-1 in vivo ^43^ and subsequent digestion by phagocytes that might result in the release of RNA from the spores intracellularly ^44,45^. We also co-cultured human PDAC samples with RJ-1 for 24 hours and found a large-scale reduction of collagen and Col-I as well as the appearance of vegetative cells inside the tumor (Fig. S10, S11). These results confirmed that RJ-1 could cause extensive collagen degradation in tumors across different species and types.

### RJ-1 mediated synthesis of INPs and induction of ferroptosis in vitro

The biochemical activity of RJ-1 to reduce iron ion (Fe^3+^) was quantified by detecting the absorbance of ferrous ion (Fe^2+^) at 562 nm after mixing with ferric citrate. As plotted in Fig. S12A, the generation of Fe^2+^ increased with the concentration of Fe^3+^ ranging from 0 to 500 µM. The absorbance of sulfide ion (S^2-^) at 665 nm was also measured after mixing RJ-1 with glutathione (GSH, a sulfur that attains high abundance within tumor tissue ^46^) or L-cysteine and a similar concentration-dependent formation of S^2-^ was observed (Fig. S12B). As expected, the production of FeS by RJ-1 increased with both the concentrations of Fe^3+^ and sulfur resources (Fig. S12C, D). Furthermore, RJ-1 could adapt to various sources of iron (ferric chloride and ferric citrate) and sulfur (GSH, L-cysteine, and Na_2_SO_3_) to synthesized FeS (Fig. S12E). Interestingly, the synthesis of FeS was observed by incubating RJ-1 in defibrinated sheep blood. It was noted that RJ-1 spores showed an adsorption effect toward Fe^3+^ (Fig. S12F), potentially due to the presence of negatively charged groups on cellular surface ^47^. To disclose the microstructure and production process of FeS, TEM was employed to analyze the culture samples of RJ-1. As displayed in Fig. 3A, high-electron-density particles, indicative of FeS, were primarily localized in the extracellular space. A portion of these particles was found to be in close proximity to or located inside the cavitated bacterial cells (Fig. 3B, C), which might be residual structures from bacterial autolysis ^48,49^. Thus, we speculated that these particles were synthesized intracellularly and released through spontaneous cell rupture after cell wall degradation, similar to the secretion of toxins A and B from *Clostridium difficile* ^49^. To verify this hypothesis, a hypertonic solution was used to block bacterial autolysis ^50^. Expectedly, numerous particles with a diverse range of sizes were observed inside the protoplast of RJ-1 (Fig. 3D). Moreover, both intracellular and extracellular particles were directly visualized with enhanced contrast in a high angle angular dark field-scanning transmission electron microscopy (HAADF-STEM) (Fig. 3E, F). As further supported by elemental analysis via energy-dispersive X-ray spectroscopy (EDS), these particles possessed both element compositions of iron and sulfur. EDS maps also revealed the enrichment of iron and sulfur in the extracellular INPs (Fig. S13A and B). These data demonstrated an efficient intracellular mineralization mediated by RJ-1 ^51^.

**Fig 3.**
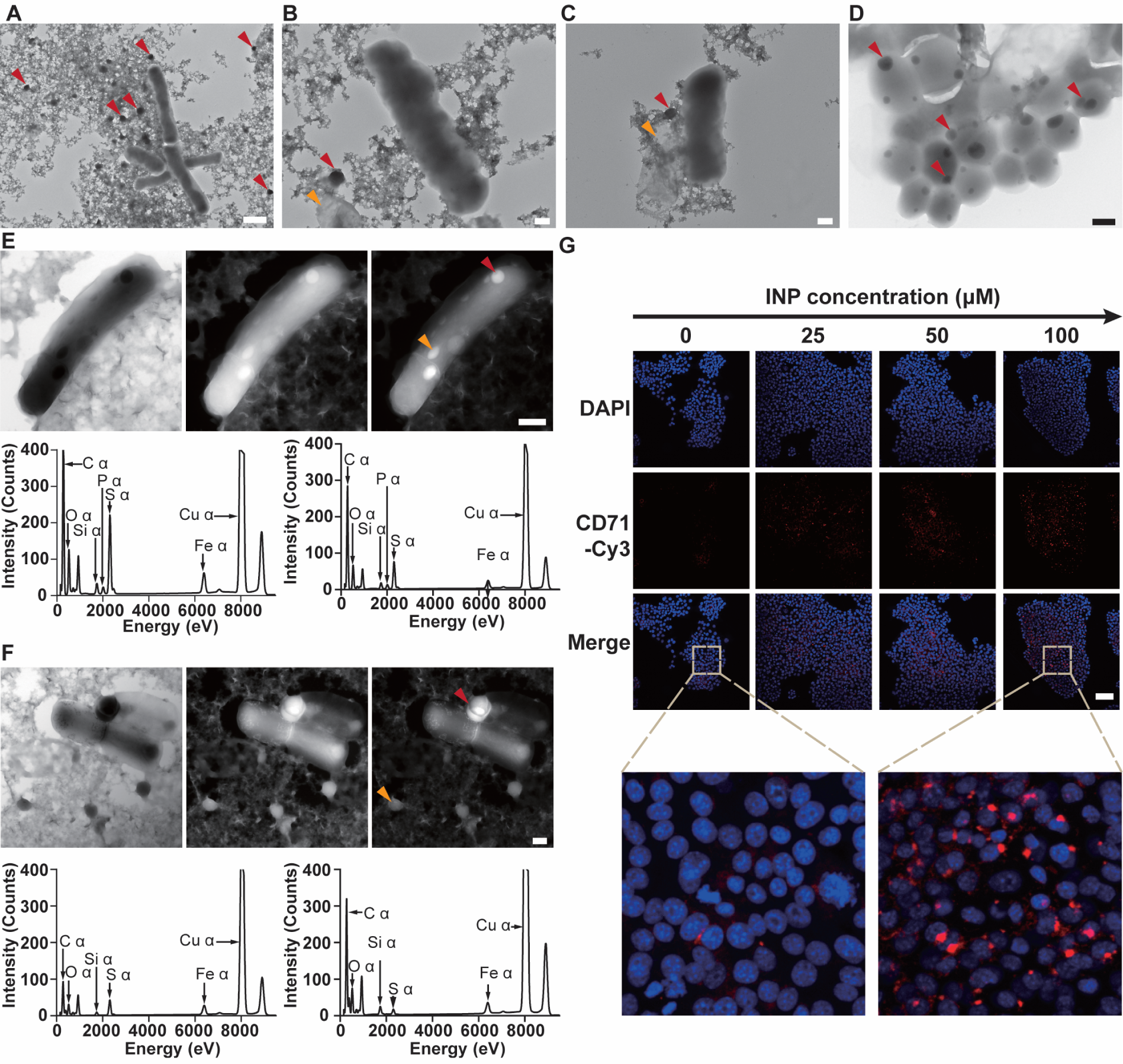
RJ-1 mediated synthesis of INPs and induction of ferroptosis in vitro. (A) A typical TEM image of INPs produced by RJ-1. Red arrows refer to INPs. Scale bar: 2 μm. TEM images of INPs (B) adjacent to and (C) inside cavitated cells. Red arrows refer to INPs and yellow arrows refer to the cavitated cell. Scale bar: 500 nm. (D) A representative TEM image of INPs in autolysis-blocked RJ-1 protoplasts. Red arrows refer to INPs. Scale bar: 500 nm. TEM imaging (bright field, dark field, and high angle angular dark field-scanning transmission electron microscopy mode from left to right) of (E) intracellularly mineralized INPs and (F) INPs released from autolytic cells and EDS analysis. The images from left to right depict bright-field, dark-field, and HAADF modes. Red arrows refer to particles corresponding to EDS results shown at left and yellow arrows refer to particles corresponding to EDS results shown at right. Scale bar: 500 nm. (G) Confocal images of 4T1 cells after treatment with a serial concentration of INPs for 4 hours. Cells were stained with CD71-Cy3 and Hoechst 33342. Scale bar: 100 μm.

Previous studies have reported that INPs can passively release ferrous ions (Fe^2+^) in the acidic environment of lysosomes upon internalization by cells, subsequently catalyzing the Fenton reaction to generate reactive oxygen species (ROS) and ultimately induce ferroptosis ^52^. Given this notion and the predominant composition of Fe^2+^ in INPs (Fig. S13C), we proposed that INPs synthesized by RJ-1 might possess ferroptosis-inducing capability. First, by methylene blue degradation assay, a strong discoloration of the solution was observed in a simulated acidic lysosomal environment (pH 5.5) by adding INPs, indicating the release of Fe^2+^, thereby triggering the Fenton reaction (Fig. S14A). Second, the catalytic activity of INPs followed a concentration-dependent response to both hydrogen peroxide (H_2_O_2_) and GSH (Fig. S14B, C), highlighting the adaptability to the tumor microenvironment ^52^. Basing on these validations, we proceeded to investigate the direct toxicity of INPs to tumor cells. As plotted in Fig. S14D, the addition of INPs significantly suppressed the proliferation of 4T1 cells (a mouse TNBC cell line), showing an even stronger effect compared to the supplementation of an equivalent amount of ammonium ferric citrate (FAC) and L-cysteine in the culture medium. As evidenced by confocal imaging, INPs induced cellular ROS production in a dose-dependent manner (Fig. S14E) and both notable cellular contraction and remarkably reduced adhesion were observed at a high particle concentration of 100 μM. Transferrin receptor 1 (CD71), a crucial receptor involved in cellular iron uptake, has been identified as a reliable marker of ferroptosis ^53^. As confirmed in Fig. 3G, by increasing the concentration of INPs, there was a significant upregulation of CD71 in 4T1 cells, particularly the formation of abundant CD71 aggregates, which closely resembled to cells treated with ferroptosis inducers, such as erastin ^53^. Briefly, INPs generated by RJ-1 could induce a unique type of cell death, namely the ferroptosis of tumor cells.

### Intratumoral synthesis of INPs and induction of tumor cell ferroptosis in vivo

Next, we examined whether RJ-1 could produce INPs in-situ after intravenous injection in TNBC-bearing mice. The tumors were collected and sectioned at 6, 12, and 18 hours post injection, respectively. Morphological structure analysis of the tumor sections revealed the occurrence of necrosis. Particularly, at 18 hours post-injection, hematoxylin-eosin (H&E) staining showed widespread presence of brown to black particles (highlighted by red arrows) in the necrotic region of the tumor (Fig. 4A), resembling hemosiderin in both color and shape ^54^. Through Gram staining, RJ-1 bacterial cells (pointed by yellow arrows) were found to be spatial proximity with these particles under both dark-field microscopy and bright-field microscopy (Fig. 4B). Elemental composition analysis of the tumor samples revealed an enlarged S^2-^ peak by X-ray photoelectron spectroscopy (Fig. 4C) and an increased total S^2-^ concentration (Fig. 4D), suggesting the presence of sulfur metabolism activity by germinated RJ-1 (Fig. S9). To verify whether these particles were RJ-1-produced INPs and corresponding INP aggregates, ultra-thin tumor tissue was sampled for TEM observation. With the help of negative staining, high-electron-density particles were observed inside bacterial vegetative cells (Fig. 4E, indicated by red arrow) and being released through autolysis (Fig. 4F and Fig. S15A). To avoid the interference of ions in the staining solution on EDS measurement, element composition analysis was also performed using unstained samples. Two distinct INP compositions were identified, one containing both iron and sulfur (Fig. 4G and Fig. S15B), and the other consisting iron alone (Fig. 4H and Fig. S15C). The latter, due to the smaller size and a more homogeneous composition, was assumed to be an intracellular breakdown product of the former ^55^. To further support the production of INPs by RJ-1, unstained sections were examined under an HAADF-STEM mode. As tumor tissue structures and bacterial structures were clearly displayed, the formation process of INPs within RJ-1 cells were well acquired. As indicated in Fig. 4I, RJ-1 exhibited the capability to form intracellular sulfur-containing particles, suggesting a preparatory step for subsequent capture of iron ions ^56^. This finding aligned with the presence of sulfur in the intracellular iron particles (Fig. 4J). Moreover, EDS analysis demonstrated the consistent element composition of INPs during in vitro culture and in the host lifecycle of RJ-1 (Fig. 3E and Fig. 4J). Given this evidence, we concluded that RJ-1 underwent the entire process of intratumoral synthesis and release of INPs in vivo.

**Fig 4.**
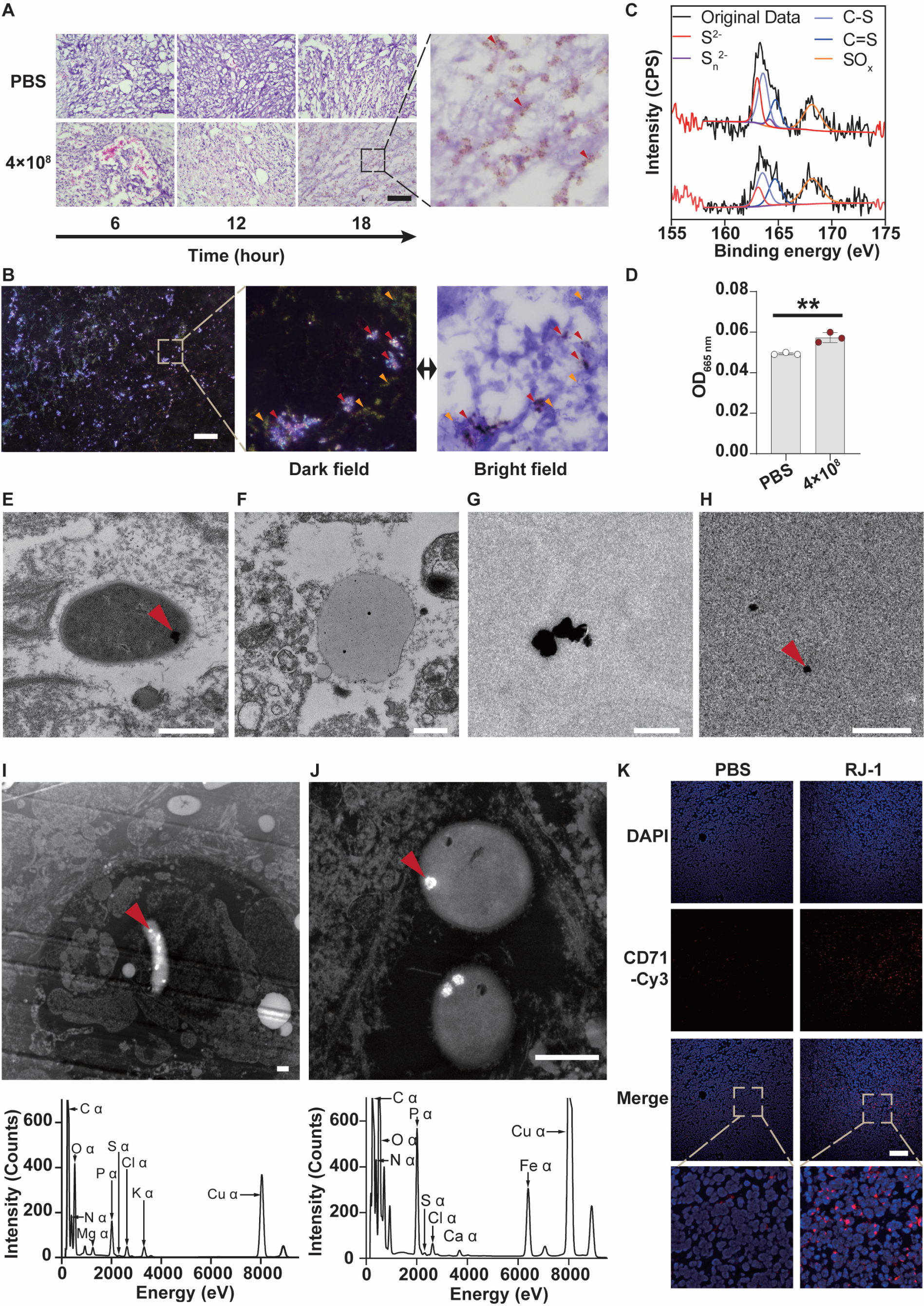
Intratumoral synthesis of INPs and induction of tumor cell ferroptosis in vivo. (A) H&E staining of tumor tissues sectioned from TNBC mice at the indicated time points after intravenous injection with 4 × 10^8^ RJ-1 spores or PBS. Red arrows refer to particle deposition. Scale bar: 100 μm. (B) Gram staining of tumor tissues sampled 18 hours post injection with 4 × 10^8^ RJ-1 spores. Images were acquired by a dark field microscopy. Red arrows refer to particle deposition and yellow arrows refer to vegetative cells. Scale bar: 100 μm. (C) X-ray photoelectron spectra of freeze-dried powder of tumor tissues collected 18 hours post injection with PBS or 4 × 10^8^ RJ-1 spores. (D) Total S^2-^ concentration in tumor tissues sampled 18 hours post injection with PBS or 4 × 10^8^ RJ-1 spores by quantifying methylene blue generated via the reaction of S^2-^ with zinc acetate, N, N-dimethyl-p-phenylenediamine, and ammonium iron sulfate. Absorbance of methylene blue was recorded at 665 nm (*n* = 3). TEM imaging of (E) vegetative cells with intracellular high-electron-density particles (red arrow), (F) high-electron-density particles released via autolysis, (G) INPs contained iron and sulfur, and (H) INPs contained iron only (red arrow pointed particle for EDS analysis) in negative-stained ultra-thin tumor sections collected 18 hours post injection with 4 × 10^8^ RJ-1 spores. Scale bar: 500 nm. HAADF-STEM imaging of vegetative cells with intracellular (I) particles containing sulfur and (J) INPs in unstained tumor ultra-thin sections obtained 18 hours post injection with 4 × 10^8^ RJ-1 spores and EDS analysis of the particles pointed by red arrows. Scale bar: 500 nm. (K) Immunofluorescence staining of tumor tissues sectioned 18 hours post injection with 4 × 10^8^ RJ-1 spores or PBS. Cells were labeled with CD71-Cy3 and DAPI. Scale bar: 100 μm. For statistical analysis of S^2-^ concentration, unpaired student’s *t* test was used to generate *p* value. **, *p* < 0.01. Error bar refer to mean value ± SEM.

In light of the in vitro induction of tumor cell ferroptosis, an intriguing question raised regarding the function of RJ-1-derived INPs within the tumor. Given the wide application of chemically synthetic iron-based nanoparticles for tumor treatment ^57^, we hypothesized that the distribution of these in-situ generated INPs in viable tumor tissue might serve as an independent cytotoxic factor for tumor cells (Fig. S16). Consistently, we observed apparent fluorescent signal corresponding to CD71 aggregates in tumor cells from tumor tissue sampled from mice dosed with RJ-1 spores at 18 hours after injection (Fig. 4K), at which time RJ-1 exhibited vigorous growth and INP production ability. TEM images suggested that tumor cells exhibited typical features of ferroptosis, including mitochondrial shrinkage, the condensation of mitochondrial membranes, and the disorganization and disappearance of mitochondrial cristae (Fig. S15D and E) ^58^. Collectively, intratumoral generation of INPs could contribute to the cytotoxic effect of RJ-1, representing an independent virulence factor named “inorganic toxin” to trigger tumor cell ferroptosis in vivo.

### Therapeutic values of RJ-1 against TNBC and PDAC

Having confirmed the capabilities of RJ-1 to degrade dense ECM and induce tumor cell ferroptosis both in vitro and in vivo, we turned our attention to explore its potential for treating cancer, particularly the ECM-dense tumors. We first conducted a dose-escalation experiment in orthotopic TNBC-bearing mice with an initial tumor of ∼200 mm^3^. Different from the doses of 10^6^ and 10^7^, a trend of delayed tumor growth was observed following a single injection dose of 10^8^ RJ-1 spores (Fig. 5A). It was worth noting that at all these doses, RJ-1 exhibited a superior tumor-targeting effect, with more than 5,000-times higher than that in the normal organs (Fig. S17A, B). Moreover, we did not observe any necrosis or living bacteria in healthy organs by H&E staining (Fig. S18A) and Gram staining (Fig. S18B), respectively. These results suggested that RJ-1 majorly germinated within the tumor, potentially suffering rapid clearance by the immune system in the periphery, indicating its satisfactory safety for in vivo application. Considering extensive tumor necrosis and rapid destruction of pre-existing collagen mesh after intravenous injection of 4 × 10^8^ spores as well as the advantage of bacterial amplification in large tumors ^59^, we then tested the tumor inhibition efficacy of RJ-1 at this dose in mice bearing TNBC with a mean volume of ∼550 mm^3^. As expected, compared to the PBS control group, significant macroscopic oncolysis occurred immediately in the mice treated with RJ-1 spores, showing substantial differences in tumor volume throughout the follow-up period (Fig. 5B).

**Fig 5.**
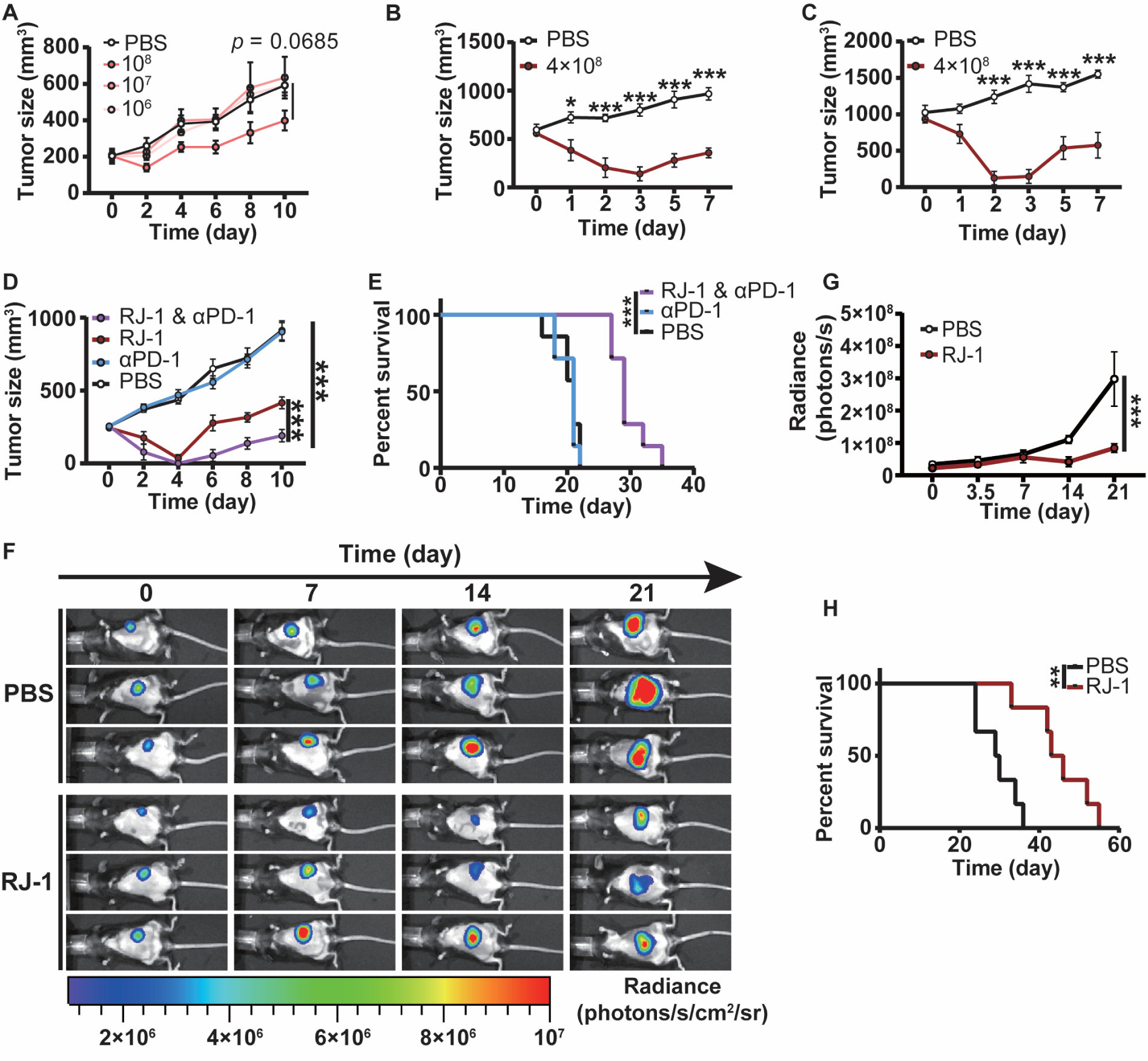
Therapeutic values of RJ-1 against TNBC and PDAC. (A) Tumor growth of orthotopic TNBC mice after treatment with the indicated doses of RJ-1 spores (*n* = 4). (B) Oncolytic effect of RJ-1 on mice with ∼550 mm^3^ orthotopic TNBC tumors before treatment (*n* = 7). (C) Oncolytic effect of RJ-1 on mice bearing ∼1000 mm^3^ giant orthotopic TNBC tumors (*n* = 7). (D) Synergistic antitumor effects of the combination of RJ-1 (4 × 10^8^ spores) and αPD-1 on orthotopic TNBC mice (*n* = 7 or 10). (E) Survival curves of orthotopic TNBC mice after different treatments (*n* = 7). (F) Typical bioluminescence images of orthotopic PDAC mice after injection with PBS or 10^9^ RJ-1 spores by IVIS (*n* = 6). (G) Tumor growth of orthotopic PDAC by measuring bioluminescence signals. (H) Survival curves of orthotopic PDAC mice after different treatments (*n* = 6). For statistical analysis of tumor growth, two-way ANOVA was used. For multiple comparison and *p* value generation, Tukey test was used in dose-escalation tumor growth experiment and Sidak test was used in other tumor growth experiments. For statistical analysis of survival, Gehan-Breslow-Wilcoxon test was used to generate *p* value. *, *p* < 0.05; **, *p* < 0.01; ***, *p* < 0.001. Error bar refer to mean value ± SEM.

The occurrence of infection-associated toxicity and cytokine storm syndrome during oncolysis may result in treatment intolerance ^60,61^. As reported, deaths have been caused upon administration of Clostridium spores to mice with tumor sizes ranging from 600 to 900 mm^3^ ^32^. To investigate whether similar risks existed during RJ-1 application, we performed intravenous injection of RJ-1 spores into mice bearing TNBC tumors with an average size of ∼1000 mm^3^. Unsurprisingly, a sharp decrease in tumor volume was observed within 72 hours, indicating significant production of necrotic material and vigorous bacterial activity (Fig. 5C). Importantly, no death was occurred as <15% body weight loss was observed to all the treated mice, even with tumor size reaching up to ∼1000 mm^3^, which was assumed to be partially attributed to mass loss caused by tumor ablation (Fig. S18C-E). Together with high treatment tolerance, extraordinary oncolytic efficacy of RJ-1 against giant tumors demonstrated its promising for future translation.

We further investigated the potential of RJ-1 in combination with existing therapies. Due to the blockage of collagen mesh to high molecular weight antibodies and the strong immunosuppressive microenvironment supported by ECM in the tumor, immunotherapy targeting programmed death-1 (PD-1) shows minimal benefits in TNBC ^62^. We speculated that by breaking down the stromal barriers and inducing tumor cell ferroptosis, RJ-1 could promote immunotherapeutic sensitivity and achieve enhanced antitumor efficacy on TNBC in combination with anti-PD-1 antibody (αPD-1). To verify this hypothesis, single-dose administration of RJ-1 spores was added into conventional αPD-1 treatment process in TNBC-bearing mice. As shown in Fig. 5D, similar to the PBS control, αPD-1 showed an ignorable therapeutic response. Strikingly, with the colonization of RJ-1, αPD-1 not only effectively inhibited tumor growth, but also achieved a significant treatment efficacy superior to that of RJ-1. We also monitored the animal survival after treatment and found that the combination therapy significantly extended the survival time of TNBC mice, by increasing the median survival from 21 days to 29 days (Fig. 5E). Note that the combination of αPD-1 and RJ-1 exhibited favorable safety as the maximum loss of body weight during treatment was limited to ∼10% (Fig. S18F).

The promising results obtained in TNBC encouraged us to evaluate the therapeutic effect of RJ-1 in orthotopic PDAC-bearing mice. Luciferase-expressing KPC 1199 cell line was used to develop the tumor model given its convenience of using luminescence to quantify tumor growth by in vivo imaging system (IVIS). As displayed in Fig. 5F, reduced bioluminescent signals of the tumors were observed in mice after RJ-1 spore administration, representing an oncolytic effect rather than merely slowing down tumor growth. Moreover, on day 21, the average bioluminescent signal of the tumors from mice treated with RJ-1 spores showed 3-times lower than that in the control group, indicating a strong therapeutic response (Fig. 5G). Accordingly, animal survival in the RJ-1 administration group was significantly elongated, with a median survival time extended by 15 days (Fig. 5H). In addition to its potency, these results also demonstrated the ability of RJ- 1 to play therapeutic effects across different tumor types. Similarly, we did not observe any animal death during this treatment and the maximum loss of body weight was in a tolerable degree of ∼10% (Fig. S18G). Overall, RJ-1 demonstrated its appealing effectiveness in treating different types of tumors with a wide range of tumor volumes.

## Discussion

We have discovered an oncolytic Clostridium strain, which expresses collagenases and produces INPs that augment its potential as a promising therapeutic agent with an attractive mechanism for combating solid tumors. Due to the presence of collagenase expression, RJ-1 exhibits the capability of degrading dense ECM both in vitro and in vivo. Notably, its ECM-degradation ability can respond to the variation in collagen concentration, functioning akin to the designed responsive synthetic biological circuits ^63^. By virtue of its nature to target tumor sites, RJ-1 is able to play a facilitating role in depleting collagen in collagen-rich tumors. Moreover, collagenases derived from RJ-1 can efficiently degrade pre-existing collagen mesh within tumors and rapidly open an accessible window for other therapeutic agents ^64^. Another characteristic of RJ-1 is its controllable production of INPs. Similar to the previously identified magnetotactic bacteria that can synthesize intracellular magnetosomes and ferrosomes ^65^, RJ-1 undergoes an intracellular mineralization process to produce INPs. Furthermore, we visualize the in-situ production of INPs in tumors through TEM observation, providing direct evidence that bacterial mineralization can serve as a source of metal particles in vivo ^66–68^. Additionally, we report the function of INPs in the interaction between RJ-1 and the host, acting as an independent virulence factor named “inorganic toxin” to induce tumor cell ferroptosis.

In the context of antitumor application, we have disclosed that RJ-1 holds a great potential for safe usage across a broad range of tumor volumes. Given the selective germination in tumor sites following systemic administration, RJ-1 can rapidly disrupt massive tumors while ensuring the survival of all experimental individuals, indicating its superior therapeutic efficacy and negligible toxicity. Markedly, intravenous injection with RJ-1 spores can induce significant tumor regression and largely prolonged survival even in visceral tumors such as PDAC without causing severe side effects. In addition to its standalone effectiveness, RJ-1 also demonstrates an appealing synergetic effect while combining with immunotherapy, highlighting its potency as a complementary approach for clinical therapies.

In summary, our work finds an oncolytic bacterial strain, with two distinct antitumor mechanisms that have been rarely reported before. RJ-1 shows encouraging prospects as monotherapy for tumor treatment. Also, as adjunctive approach, RJ-1 exhibits a broad effective dosage regimen and an extended time window of synergy. By applying RJ-1 as a chassis for synthetic biology, collagen degradation and ferroptosis induction can be incorporated as fundamental functionalities into existing engineering strategies. These discoveries pave an avenue for developing advanced bacteria-based therapeutics for the treatment of cancer and beyond.

### Methods Cell culture

NIH/3T3 cells were obtained from Chinese Academy of Sciences Collection Committee cell bank (Shanghai, China). 4T1 cells were obtained from American Type Culture Collection. The primary tumor cell line KPC1199, originating from KPC pancreatic ductal adenocarcinoma mouse model (LSL-KrasG12D/+; LSL-Trp53^R172H/+^; Pdx-1-Cre, on C57BL/6 background), was generously provided by Dr. Yiyi Liang. The KPC1199 cell line was initially kindly shared by Prof. Jing Xue from Renji Hospital (Shanghai, China).

To establish luciferase-expressing KPC1199 cell line, lentiviral particles containing the luciferase gene along with a blasticidin S-resistance gene, were used to transfect KPC1199 cells (MOI=10). Following transfection, the cells were subjected to continuous selection with blasticidin S (100 μg/ml) for a period of at least two weeks to obtain a stable transfected luciferase-expressing KPC1199 cell line.

NIH/3T3, KPC1199, and luciferase-expressing KPC1199 cells were cultured in Dulbecco’s Modified Eagle Medium (DMEM) supplemented with 10% fetal bovine serum (FBS), 100 μg/mL streptomycin and 100 units/mL penicillin. 4T1 cells were cultured in Roswell Park Memorial Institute 1640 medium supplemented with 10% FBS, 100 μg/mL streptomycin, and 100 units/mL penicillin. All cultures were maintained at 37 °C with a 5% CO_2_ atmosphere.

### Bacterial culture

For acquirement of bacteria at logarithmic growth state, a single colony of *Clostridium bifermentans* RJ-1 (RJ-1) was picked and cultured anaerobically in brain heart infusion (BHI) medium at 37 °C for 72 hours. On the following day, the RJ-1 culture was sub-cultured into fresh medium at a 1:100 ratio and continued to grow for 4 hours before collection and storage at 4 °C in an anaerobic bag for subsequent use. Other bacteria were cultured in Lysogeny broth (LB) medium. The bacterial growth in liquid culture was determined by measuring the absorbance at a wavelength of 600 nm.

To harvest spores, RJ-1 was cultured in BHI medium under an anaerobic condition for one week. The culture was then centrifuged at 13,000 *g* for 3 minutes to collect bacteria and spores, followed by two washes with phosphate-buffered saline (PBS). Different concentrations of sucrose solutions, including 30%, 40%, and 50%, were prepared for density gradient centrifugation. The sucrose solutions were layered in 15 ml centrifuge tubes from high to low concentration, and then the PBS-suspended clostridial spores and bacteria mixture was layered on top. The tubes were centrifuged at 800 g for 15 minutes, making spores enrich in 30% and 40% layers, and bacteria in 50% layer and the pellet. Isolated vegetative cells and spores were imaged under dark-field microscopy to distinguish their morphology.

For bacterial enumeration, 50 μL of the bacterial suspension was spread on clostridial differential agar containing 0.4 g/L D-cycloserine in 10 cm dishes. The plates were incubated at 37 °C under the anaerobic condition for 12-16 hours, and the number of bacterial colonies on the plates was counted. Black bacterial colonies of RJ-1, representing ferrous sulfide production on clostridial differential agar, were photographed after 24 hours of incubation.

To characterize the hemolytic ability of RJ-1, 50 μL of the bacterial suspension was spread on Columbia blood agar plates, and hemolytic zones were photographed after 24 hours of incubation. To characterize the swarming motility and autolysis of RJ-1, 5 μL bacterial culture in logarithmic phase was spotted in the center of agar plates, and colony morphology was recorded after 7 and 10 days of growth.

To create skim milk agar plates, 3% sterilized skim milk was added into suitable agar plates for different bacteria. For example, *Enterococcus faecalis*, *Bacillus cereus*, *Escherichia coli* Nissle-1917, and *Salmonella typhimurium* VNP20009 (VNP20009) were cultured in LB plates containing skim milk, while RJ-1 was cultured on clostridial differential agar with skim milk.

### Detection of sulfide ion and ferrous ion

For sulfur metabolism, different concentrations of glutathione (GSH) or L-cysteine (L-cys) were added to BHI solution with 2 mM zinc chloride. Spores were added to 1 mL of the culture medium and anaerobically cultured for 24 hours. The culture was then centrifuged at 10,000 *g* for 10 minutes, and the pellet was washed once with 5 M sodium hydroxide and once with water. Finally, an equal volume of 1 M hydrochloric acid solution containing 3 mM N, N-dimethyl-p-phenylenediamine sulfate and 6.6 mM ferric chloride was added. The reaction was carried out at room temperature for 30 minutes, and the absorbance at 665 nm of the supernatant was measured. The sulfur ion measurement in tumor tissue harvested at 18 hours post-treatment was performed by using a hydrogen sulfide assay kit.

In BHI culture medium, different concentrations of ammonium ferric citrate (FAC) were added, and the culture was incubated anaerobically with or without spores. After 24 hours, a 2.5 mM ferrozine solution (diluted 1/10 in the culture medium) was added, and the mixture was allowed to develop color at room temperature for 20 minutes. The absorbance was measured at 562 nm.

### Ferric ion adsorption assay

10^8^ RJ-1 spores or equivalent volume of distilled deionized water were added to a 100 μM ferric chloride solution and incubated at 37 °C for 2 hours. After incubation, the mixture was centrifuged at 13,000 *g* for 3 minutes to remove the precipitate. The supernatant was collected, and 1/10 volume of 2.8 M hydroxylamine hydrochloride solution and 1/10 volume of 2 mM ferrozine solution were added. The mixture was thoroughly mixed and allowed to stand at room temperature for 20 minutes. The absorbance at 562 nm was measured to determine the extent of iron ion adsorption.

### Production and collection of iron-containing particles (INPs)

RJ-1 was cultured in BHI broth containing 2 mM FAC and 6 mM L-cys for 12 hours. The culture was then centrifuged at 13,000 *g* for 10 minutes, and the pellet was transferred to a 0.33 M sucrose solution overnight to promote cell wall degradation. Subsequently, the cells were treated with 1% Triton X-100 to disrupt the protoplasts and release the intracellular particles.

To verify the contribution of different sulfur and iron sources to INPs production, various substrates containing 2 mM iron and 6 mM sulfur elements were added to BHI medium. RJ-1 spores were inoculated, and the cultures were anaerobically incubated at 37 °C for 24 hours.

To investigate if blood could serve as an iron source for INP production, defibrinated sheep blood was mixed with BHI medium in a 1:4 volume ratio. RJ-1 spores were added to the mixture and incubated for 24 hours under a condition of 6 mM L-cys.

To assess the iron and sulfur source concentration-dependent production characteristics of INPs, different concentrations of FAC and GSH were added to BHI solution, and bacterial spores were incubated for 24 hours. INPs were collected and treated with an equal volume of 2 mM ferrozine hydrochloride solution. After a 2-minute reaction at room temperature, a small amount of sodium hydroxide solution was added to adjust the pH to 5. After centrifugation to remove the precipitate, the absorbance of the solution was measured at 562 nm.

### Degradation of collagen by RJ-1 in vitro

For acid-soluble collagen degradation, 10^8^ colony forming unit/ml (CFU/mL) of RJ-1 spores or VNP20009 bacteria were mixed with 0.5% BHI agar and poured into bacterial culture dishes. The cultures were anaerobically incubated for 24 hours to form a bacterial lawn. Mouse tail collagen was dissolved in 0.02 M HCl and neutralized to a neutral pH with 0.25 M Tris(hydroxymethyl)aminomethane (Tris) and 4 M NaCl. A 3 mL aliquot of the neutralized collagen solution was placed in each culture dish to form a gel. Circular bacterial lawns were inverted and placed in the center of the gel, and collagen degradation around the bacterial lawn was monitored.

For the characterization of cell-derived collagen degradation, NIH/3T3 cells were seeded at 10^4^ cells per well in a 96-well plate and cultured in 100 μL of complete DMEM medium (DMEM with 10% FBS) for 24 hours. After 24 hours, the culture medium was removed, and the cells were washed three times with PBS. Subsequently, 100 μL of 1% Triton X-100 in ultrapure water was added to each well to lyse the cell monolayer, and the plate was incubated at 37 °C for 30 minutes. The Triton X-100 solution was removed, and the cells were washed three times with ultrapure water. Next, 100 μL of BHI medium or BHI bacterial culture liquid, grown from RJ-1 single colony culture for 72 hours to the plateau phase, was added to each well. The plate was incubated under an anaerobic condition for 12 hours. After incubation, the liquid was removed, and the cells were washed three times with ultrapure water. Each well was stained with 100 μL of picrosirius red solution for 30 minutes. After 30 minutes, the picrosirius red solution was removed, and the wells were soaked with 0.5% acetic acid solution for three minutes and three times. Subsequently, the wells were washed three times with ultrapure water. The degradation was visualized using an upright microscope. The picrosirius red was dissolved using 100 μL of 0.5% sodium hydroxide solution, and the absorbance was measured at 540 nm to quantify the extent of degradation.

For the characterization of collagen degradation ratio, gelatin at a concentration of 20 mg/mL was mixed with BHI medium at a 1:9 ratio, or an equal volume of PBS was used as a substitute. RJ-1 bacterial liquid grown for 72 hours was transferred at a 1:100 ratio into fresh medium containing or lacking gelatin. Bacterial liquid was collected at different time points, and the concentration of hydroxyproline (HYP) was measured. The total HYP concentration in liquid culture medium was determined using the Hydroxyproline Assay Kit (NJJCBIO, A030-3-1). And the HYP content in BHI powder and solid gelatin was determined using the same kit following acid hydrolysis.

### Detection of reactive oxygen species (ROS) production

To investigate the generation of ROS by INPs in a simulated lysosomal environment, 100 μM INPs were added to a sodium dihydrogen phosphate-citrate buffer with different pH containing 10 μg/mL methylene blue. The mixture was incubated at 37 °C for 4 hours, and the absorbance at 665 nm in the supernatant was measured.

To assess the dependency of INP-induced Fenton reaction on GSH and hydrogen peroxide concentrations, different concentrations of GSH and 2 mM hydrogen peroxide, or varying concentrations of hydrogen peroxide and 5 mM GSH, were added to a sodium dihydrogen phosphate-sodium citrate buffer (pH 5.5) containing 10 μg/mL methylene blue. Following incubation at 37 °C for 4 hours with 100 μM INPs, the absorbance of the supernatant was measured at 665 nm.

To verify whether INPs induce intracellular ROS production, different concentrations of INPs were added to confocal dishes containing 4T1 cells. The cells were incubated for 12 hours in the cell culture environment, and the intracellular ROS production was detected using the Reactive Oxygen Species Assay Kit (Beyotime, S0033S) with the 2’,7’-dichlorodihydrofluorescein diacetate (DCFH-DA) probe.

### SDS-PAGE and zymography

To investigate the presence of collagenases and determine their main molecular weights in bacterial secreted proteins, RJ-1 bacterial culture in BHI medium was incubated for 72 hours. After incubation, the culture was subjected to centrifugation at 13,000 *g* for 10 minutes to collect the supernatant. The supernatant was then filtered through a 0.45 μm membrane and ammonium sulfate was added to achieve a 50% mass fraction. The mixture was stirred on ice for 30 minutes and subsequently centrifuged at 12,000 *g* for 10 minutes to pellet the proteins in 1 L of supernatant. The proteins were then concentrated to 100 times their original concentration.

For protein electrophoresis, a 10% SDS-PAGE gel was used. Alternatively, for zymography experiments, 1 mg/mL of gelatin was incorporated into the separating gel during SDS-PAGE gel polymerization. The concentrated proteins were diluted in a 2-fold gradient four times, and 10 μL of each dilution was loaded onto either the zymography gel or the regular SDS-PAGE gel. SDS-PAGE gels were prepared using 10% Non-Closure SDS-PAGE Color Preparation kit (Sangon Biotech, C681102).

For SDS-PAGE, the gel was stained with 0.5% Coomassie Brilliant Blue R250 (dissolved in 40% methanol and 10% acetic acid) at room temperature for 30 minutes. Subsequently, the gel was destained multiple times with a methanol-acetic acid solution until clear bands were observed. For zymography analysis, gel blocks were washed four times for 15 minutes each with wash buffer (2.5% Triton X-100, 50 mM Tris-HCl, 10 mM CaCl_2_, 1 μM ZnCl_2_, and 50 mM NaCl, pH 7.6). Afterward, the blocks were washed twice for 15 minutes each with incubation buffer (50 mM Tris-HCl, 5 mM CaCl_2_, 1 μM ZnCl_2_, 50 mM NaCl, and 0.05% Brij35, pH 7.6). The gel blocks were then incubated at 37 °C for 24 hours in the incubation buffer. After incubation, the gel was washed three times with deionized water, stained with Coomassie Brilliant Blue, and destained until clear bands were visible, followed by photography. SDS-PAGE gels were prepared using 10% Non-Closure SDS-PAGE Color Preparation kit (Sangon Biotech, C681102).

For assessment of changes in total collagenase activity in tumor, following an 18-hour treatment, tumor samples from mice were collected and subjected to lysis and grinding in Radioimmunoprecipitation assay buffer (RIPA) containing a protease inhibitor cocktail (Selleck, B14001). The lysate was then centrifuged at 10,000 *g* for 10 minutes, and the supernatant containing proteins was collected. A total of 25 μg of protein was used for SDS-PAGE and zymography experiments. SDS-PAGE gels were prepared using 4-15% precast SDS PAGE gel (Shanghai Wansheng Haotian Biotechnology Co., Ltd, PSG2001-415F) and gels for zymography were prepared using 10% Non-Closure SDS-PAGE Color Preparation kit (Sangon Biotech, C681102).

### Western blot

To detect type I collagen in tumors, mice tumor samples were collected on the tenth day after treatment. The samples were lysed and homogenized in RIPA lysis buffer containing a protease inhibitor cocktail. After centrifugation at 10,000 *g* for 10 minutes, the supernatant was collected to obtain the protein. An amount of 40 mg of protein from each sample was loaded onto the gel for electrophoresis and subsequently transferred to a polyvinylidene difluoride membrane using Trans-blot Turbo. Rabbit Anti-Col-1A1 Polyclonal Antibody (Boster, BA0325) was used for detecting type I collagen, while a β-Actin Rabbit Monoclonal Antibody (Abclonal, AC038) was used to detect β-Actin as a loading control. HRP Goat Anti-Rabbit IgG (H+L) antibody (Abclonal, AS014) was applied as a secondary antibody. Chemiluminescence analysis was performed using the StarSignal Plus Chemiluminescent Assay Kit (Genstar, E170-01).

### Histochemistry

Tissues were collected and fixed in 4% paraformaldehyde, followed by embedding in either paraffin or optimal cutting temperature compound (OCT) for subsequent paraffin or frozen sectioning. For hematoxylin-eosin (H&E) staining, sections were hydrated and stained with hematoxylin, followed by differentiation in acid alcohol and counterstaining with eosin. To perform picrosirius red staining, hydrated sections were stained with a solution of picrosirius red dissolved in saturated picric acid. Gram staining was carried out using ammonium oxalate crystal violet solution, 1% iodine solution, and safranin solution. All sections were mounted with neutral mounting medium and imaged under brightfield, darkfield, or polarized light modes.

Statistical analysis of the area fraction of picrosirius red staining was performed by comparing 14 randomly selected fields of necrotic regions, perinecrotic regions, and non-necrotic regions.

### Immunofluorescence and fluorescent in situ hybridization

To detect the iron death marker CD71 in tumor tissue sections, antigen retrieval was firstly performed by using a high-temperature antigen retrieval solution. The sections were then blocked with goat serum for 30 minutes. The primary antibody was added and incubated overnight in a humidified dark chamber, followed by incubation with a fluorescent secondary antibody at room temperature for 1 hour. Finally, the sections were mounted with antifade mounting medium with 2-(4-amidinophenyl)-6-indolecarbamidine dihydrochloride (DAPI) (Beyotime, P0131). Anti- CD71 recombinant rabbit monoclonal antibody (HUABIO, JF0956) was used as primary antibody and Cy3 conjugated goat anti-rabbit IgG (H+L) (Abclonal, AS007) was used as secondary antibody.

For the staining of type I collagen on pancreatic ductal adenocarcinoma (PDAC) tumor tissue, paraffin-embedded sections were collected at different time points after drug administration. Antigen retrieval was performed using Proteinase K (Beyotime, ST532). Initially, the Clit135- FAM probe was incubated overnight at 37 ℃ in an incubation buffer consisting of 20 mM Tris-HCl (pH 7.2), 0.1% sodium dodecyl sulfate (SDS), 0.9 M NaCl, 20% formamide, and 10% sodium dextran sulfate with a probe concentration of 10 ng/μL. After incubation, the sections were washed twice at 37 ℃ using the washing buffer containing 20 mM Tris-HCl (pH 7.2), 0.1% SDS, and 0.145 M NaCl. Subsequently, the sections were blocked with goat serum before the primary antibody was added, and incubated overnight in a humidified dark chamber, followed by incubation with a fluorescent secondary antibody at room temperature for 1 hour. Finally, the sections were mounted with antifade mounting medium with DAPI (Beyotime, P0131). Anti-Collagen I/Col-1A1 rabbit monoclonal antibody (Abclonal, A22089) was used as primary antibody and ABflo® 647-conjugated goat anti-rabbit IgG (H+L) (Abclonal, AS060) was used as secondary antibody.

For immunofluorescence staining of CD71 on cells, 4T1 cells were treated with different concentrations of INPs (corresponding to different total iron concentrations) in confocal dishes, and thoroughly washed with PBS for three times before being fixed with 4% paraformaldehyde for 1 hour. Then, they were blocked with 1% bovine serum albumin (BSA) for 30 minutes, and incubated with the primary antibody and fluorescent secondary antibody. Finally, Hoechst 33342 was used to stain nucleus for 10 minutes. Anti-CD71 rabbit monoclonal antibody was used as primary antibody and Cy3 conjugated rabbit anti-goat IgG (H+L) was used as secondary antibody. The background fluorescence was eliminated during the staining process using a tissue autofluorescence quencher (Servicebio, G1221).

### Tumor spheroid formation

Tumor spheroids were generated by using the hanging drop method. A solution of 0.24% methylcellulose dissolved in full culture medium (DMEM containing 10% FBS) was prepared. In each hanging drop, 20 μL of the 0.24% methylcellulose-culture medium solution was pipetted onto the lid of a 100 mm dish. This solution contained a total of 20,000 cells, consisting of 4T1 to NIH/3T3 cells at the ratio of 2:1. The lids with the hanging drops were then inverted over dishes containing 10 mL of phosphate buffer solution. Subsequently, the hanging drop cultures were incubated under standard culture condition (5% CO_2_, 37 °C) for 4 days. After the incubation period, the tumor spheroids formed in the drops were carefully harvested for subsequent experiments.

### Liposome preparation

Liposomes containing 95% dipalmitoyl phosphatidylcholine (DPPC) and 5% distearoyl phosphoethanolamine-polyethylene glycol (DSPE-PEG) were prepared via rotary evaporation. The liposomes were loaded with 1,1’-dioctadecyl-3,3,3’,3’-tetramethylindocarbocyanine perchlorate probe at a ratio of 0.1%. The lipids were extruded using consecutive 400 nm and 200 nm polycarbonate membranes to obtain uniformly sized liposomes. The extrusion process was conducted at a temperature above 50 ℃, and a total of 10 extrusion cycles were performed using back-and-forth extrusion. The resulting nanoscale liposomes were stored at 4 ℃ for further use.

### Co-culture of tumor spheroids and bacteria

For the characterization of tumor spheroid collagen degradation, 50 μL of 1% agar was poured into each well of a 96-well plate to prevent cell adhesion to the bottom. Tumor spheroids were transferred into the wells, followed by the addition of 100 μL of DMEM culture medium containing 2.5% BHI. Subsequently, 10^7^ RJ-1 spores were added to each well. An equal volume of PBS was used as the control. After incubation at 37 °C in a 1% low-oxygen environment for 12 hours, the spheroids were removed, fixed with 4% paraformaldehyde and subjected to permeabilization and immunostaining steps. The primary antibody and secondary antibody were used for type I collagen staining and Hoechst 33342 was used for nuclei staining. After sufficient transparency with a 60% glycerol and 2.5 M fructose solution, these spheroids were imaged by confocal microscopy.

In penetration experiment, 50 μL of 1% agarose was poured into 96-well plates to prevent tumor spheroids from adhering to the bottom. Tumor spheroids were transferred into the wells with 100 μL of DMEM medium containing 2.5% BHI. A number of 10^7^ RJ-1 spores were added into each well. An equal volume of PBS was added as a control. The co-cultures were then incubated at 37 ℃ under 1% hypoxic conditions for 16 hours. Subsequently, the tumor spheroids were removed and exposed to either 10 μg/mL Hoechst or 50 μg/mL 1,1’-dioctadecyl-3,3,3’,3’- tetramethylindocarbocyanine perchlorate-labeled liposomes for 1 hour. After three washes with PBS, the tumor spheroids were fully clarified using a solution with 60% glycerol and 2.5 M fructose before being observed by confocal microscopy.

In the oncolytic assay, RJ-1 spores or VNP20009 bacteria at a concentration of 10^8^ CFU/mL were mixed with 1% BHI agar and added to a 96-well plate at a volume of 100 μL per well. An equal amount of agar without spores was added into control wells. Then, the tumor spheroids were cultured in the wells, with 100 μL of DMEM complete culture medium, at 37 ℃ under 1% hypoxic conditions. Tumor spheroid morphology was captured at different time points (0 hours, 12 hours, and 24 hours).

### Cell viability assay

4T1 cells were seeded at a density of 10,000 cells per well in a 96-well plate. The cells were then treated with different compounds, including PBS, 100 μM FAC with 300 μM L-cys, or INPs containing 100 μM iron. The treated cells were continuously monitored using Incucyte S3 for 24 hours. Cell viability was assessed based on the measurement of the total cell area.

### Scanning electron microscopy of collagen mesh

After different hours of RJ-1 spores or PBS administration, tumor tissues were collected and immersed in a 1% SDS aqueous solution to remove cells. The remaining collagen meshes were fixed in a 2.5% glutaraldehyde solution after washing with water. Subsequently, the fixed tissues were subjected to a dehydration process using a series of ethanol concentrations: 30%, 50%, 70%, 90%, 95%, and 100%. After replacing ethanol with tert-butanol, the samples were cured at 4 ℃ and then vacuum-dried at room temperature. The obtained samples were then imaged using scanning electron microscope to visualize the surface and cross-sectional structures.

### Transmission electron microscopy (TEM)

To visualize the vegetative cell and spore morphology of RJ-1, 5 µL of vegetative cell suspension or spore suspension was pipetted onto a copper grid with carbon film and allowed to settle for twenty minutes. Excess liquid was removed, and the grids were air-dried to prepare TEM samples for subsequent imaging.

For visualization of INPs production in vitro, RJ-1 was cultured in BHI medium containing 2 mM FAC and 6 mM L-cys for 12 hours. After incubation, 5 μL of the cultured liquid was deposited onto a copper grid with carbon film and allowed to settle for twenty minutes. Excess liquid was removed, and the grids were air-dried to prepare TEM samples for subsequent imaging. To observe protoplasts, BHI medium was mixed with 0.35 M sucrose at a ratio of 1:9, and 2 mM FAC and 6 mM L-cys were added to the culture system. The mixture was then cultured for 12 hours. After incubation, 5 μL of the cultured liquid was deposited onto a copper grid with carbon film and allowed to settle for twenty minutes. Excess liquid was removed, and the grids were air-dried to prepare TEM samples for subsequent imaging.

Tumor tissues were collected after RJ-1 intravenous administration for 18 hours. The tumor sample was fixed in glutaraldehyde solution and post-fixed in osmium tetroxide. Dehydration of the tissue was carried out using a series of graded ethanol to replace water in the sample. Samples were then infiltrated and embedded with resin. Ultra-thin sections were gained by using an ultramicrotome, collected on copper grids, and stained by uranyl acetate.

Images were captured under bright filed mode, dark field mode or high angle angular dark field-scanning transmission electron microscopy mode, and energy-dispersive X-ray spectroscopy (EDS) and EDS map was carried out for analyzing high electron density particles.

### Atomic force microscopy

Frozen cryo-sectioned tissue samples were thawed and thoroughly washed with PBS to eliminate residual OCT. Subsequently, the samples were imaged at room temperature using the force spectroscopy mode of AFM. DNP-10 with a triangular tip radius of 0.6 μm was employed for imaging. The spring constant of the cantilever was 0.35 N/m. During imaging, the force-indentation curves were generated with a velocity of 20 μm/s and a maximum force of 8 nN. To derive a topography and elastic modulus (E-modulus) image from each force map, the approach part of every force-indentation curve was fitted using the Hertz model. To compare the E-moduli between the control and tumor tissues, four force maps, each consisting of 16 × 16 force-indentation curves and covering a scan area of 25 × 25 μm^2^, were recorded at various locations within each tissue sample. From each force map, an average E-modulus was calculated.

### **X-** ray photoelectron spectroscopy

Tumor tissue was collected after RJ-1 or PBS administration for 18 hours, rapidly frozen in liquid nitrogen, and then cryogenically ground. The resulting powder was freeze-dried to prepare the sample for XPS analysis. Calibration of the binding energies was performed using the carbon 1 s peak at 284.8 eV. Fitting of the spectra was performed using CasaXPS.

To determine the valence state distribution of iron, INPs were collected, vacuum dried, and subjected to XPS analysis. Calibration of the binding energies was performed using the carbon 1 s peak at 284.8 eV. Fitting of the spectra was performed using CasaXPS.

### Animal model establishment and in vivo imaging

To establish an orthotopic triple-negative breast cancer (TNBC) model, 6-8-week female Balb/C mice were used. 4T1 and NIH/3T3 cells were mixed in a 2:1 ratio and a total of 100,000 cells were injected into the right third mammary fat pad of the mice with Matrigel (corning, 356234). Once the tumors reached a certain size, treatment could be conducted. For intratumoral injection, when the tumor volume reached approximately 200 mm^3^, 10^6^ bacteria were resuspended in 20 μL PBS and injected into the tumor. Tumor samples were collected on the tenth day after injection. For the dose escalation experiment, when the tumor volume reached around 200 mm^3^, mice were divided into four groups, consisting of 4 mice in each group. 10^6^, 10^7^, and 10^8^ RJ-1 spores were resuspended in 100 μL PBS and injected via the tail vein. Mouse samples were obtained on the tenth day for intratumoral collagen detection and organ safety assessment. The tumor HYP content was determined using Hydroxyproline Content Assay Kit (Beijing Boxbio Science & Technology Co.,Ltd, AKAM017C), and normalization was performed based on the tumor mass weighed on an analytical balance (METTLER TOLEDO, ME104E/02). For tumors at the average sizes of approximately 550 mm^3^ and 950 mm^3^, 4 × 10^8^ RJ-1 spores were resuspended in 100 μL PBS and injected intravenously, with 7 mice per group. For combination therapy, when the tumor volume reached around 250 mm^3^, 4 × 10^8^ RJ-1 spores were resuspended in 100 μL of PBS and injected intravenously on day 1, and 200 μg of anti-PD-1 antibodies (αPD-1, Bio X Cell, RMP1-14) was injected intraperitoneally on day 0, 3, and 6. For combination therapy experiment, 7 mice were used in PBS, αPD-1 and αPD-1 & RJ-1 groups, and 10 mice were used in RJ-1 group. For all

TNBC-bearing mice, tumor volume was estimated by the following formula of 0.5 × length × (width)^2^. Once tumor growth reached or exceeded 2000 mm^3^, mice were considered to have reached the endpoint and were euthanized.

To establish a PDAC model, 10,000 luciferase-expressing KPC1199 cells were resuspended in 25 μL of PBS and injected into the pancreatic tail of C57BL/6 mice. Tumor growth was monitored by injecting 3 mg of D-Luciferin potassium salt intraperitoneally and detecting bioluminescence 15 minutes later. Once tumor radiance reaching 10^6^ (photons/s), 10^9^ bacterial spores resuspended in 500 μL of PBS or equivalent volume of PBS was injected intraperitoneally into PDAC-bearing mice. Tumor samples were collected on day 1, 2, 7, and 14 after injection.

All animals used were purchased from SPF (Beijing) biotechnology co., Ltd. Animal experiments were performed under the guidelines evaluated and approved by the ethics committee of Institutional Animal Care and Use Committee of Shanghai Jiao Tong University (A2023017).

### Tumor penetration analysis of fluorescein isothiocyanate conjugated dextran (dextran-FITC)

For preparation of dextran-FITC: a solution was prepared by dissolving 1.1 g of dextran (70 kDa) in 10 mL of DMSO containing a few drops of pyridine. Next, 0.1 g of FITC was added followed by the incorporation of 20 mg dibutyltin dilaurate (1.066 mg/µL). The mixture was then incubated at 95 °C for 2 hours. Unbound FITC was removed by multiple ethanol precipitations. Vacuum drying is carried out to obtain the final product.

For tumor penetration analysis in vivo: mice bearing orthotopic TNBC were injected with dextran-FITC intravenously at 250 μg/g dose. One-hour post-injection, mice were anesthetized with isoflurane. Skin overlying the tumor was then carefully dissected, and the distribution of dextran-FITC within the tumor region was observed using an Optiscan in vivo confocal microscopy system.

### Tumor opening assay

Mice with a volume of 200 mm³ TNBC accepted 10^6^ spores or PBS intravenous injection, and tumors were harvested on the tenth day after injection. The tumors were cut along their longest axis, with a cut depth equal to 80% of the shortest axis length, then were allowed to relax in PBS for 10 minutes, and the opening width was measured in the middle of the incision. Normalization was performed based on the tumor diameter in the vertical direction of the cut.

### Co-culture of human PDAC sample with bacteria

The protocols were approved by the approval of Human Ethics Committee of Renji Hospital affiliated to Shanghai Jiao Tong University School of Medicine (KY2023-020-B). Human PDAC samples were collected from clinical sources and sliced into 1 mm thick sections. Co-cultivation was performed using the gas-liquid interface method. Briefly, a gelatin sponge was placed at the bottom of a 6-well plate, and complete culture medium (Dulbecco’s Modified Eagle Medium/Nutrient Mixture F-12 with 10% FBS) containing 2.5% BHI was added to each well, ensuring that the liquid level just covered the gelatin sponge. The human PDAC samples were then placed on the gelatin sponge. Subsequently, 4 × 10^8^ RJ-1 spores were added to the culture medium, and the co-cultures were maintained under cell culture conditions (5% CO_2_, 37 °C) for 24 hours for subsequent sectioning and analysis. An equivalent volume of PBS was added as a control.

### Genome sequencing and 16S ribosomal RNA (rRNA) phylogenetic tree generation

RJ-1 genome sequencing and analysis were conducted by Sangon Biotech. In brief, genome DNA was extracted from cultured RJ-1. Sequencing was performed using Illumina Hiseq™ to acquire raw data. Quality assessment of the sequencing raw data was conducted using FastQC. The Illumina sequencing data was subjected to quality trimming using Trimmomatic to obtain refined and accurate usable data. Genome assembly was accomplished using SPAdes for assembling the second-generation sequencing data. Gap filling of contigs was achieved with GapFiller. Sequence correction was carried out using PrInSeS-G to rectify clipping errors and small fragment insertions or deletions introduced during the assembly process. For genome analysis, gene elements including genes, tRNAs, and rRNAs were predicted using Prokka. The predicted genes were then subjected to functional annotation by comparing their protein sequences against various databases such as CDD, KOG, COG, and VFDB using NCBI Blast. Additionally, the predicted 16S rRNA sequences were aligned with the NCBI 16S database using NCBI Blast to ascertain homologous strains, and representative strains from the 11 bacterial species with the highest homology to RJ-1 were utilized for the construction of a phylogenetic tree.

### Statistical analysis

The data obtained in this study underwent rigorous statistical analysis using appropriate methods to determine the significance of observed differences. For multiple group comparisons, ordinary one-way ANOVA with Tukey’s multiple comparison test was utilized to assess differences among three or more groups. Additionally, two-way ANOVA with Tukey’s post hoc test or Sidak test was employed to investigate the effects of two independent factors on the experimental outcomes. For statistical analysis of survival, Gehan-Breslow-Wilcoxon test was used to generate p value. Furthermore, comparisons between two groups were performed using student’s *t* test, which provided a robust assessment of mean differences, considering the variance within the groups. The significance level was set at *p* < 0.05. Results were considered statistically significant when *p* values were below the chosen threshold.

### Data availability

All data needed to evaluate the conclusions in the paper are present in the paper and/or the Supplementary Information. Data is available from the authors on request.

## Acknowledgments

This work was financially supported by the National Key Research and Development Program of China (2021YFA0909400), the National Natural Science Foundation of China (22305152, 32101218, 22375127), the Explorer Program of the Science and Technology Commission of Shanghai Municipality (21TS1400400), the Talent Promotion Program of Renji Hospital (RJTJ22-RC-002, RJTJ23-RC-010), the Innovative Research Team of High-Level Local Universities in Shanghai (SHSMU-ZDCX20210900, SHSMU-ZDCX20210700), the Foundation of National Infrastructures for Translational Medicine (Shanghai) (TMSK-2021-123, TMSK-2021-119), and the Shanghai Municipal Education Commission-Gaofeng Clinical Medicine Grant Support (20181704, 20191820).

## Author contributions

J.L. supervised the project. J.L. and F.Y. conceived and designed the experiments. F.Y., L.M., H.C., X.W., M.Z. and Z.C. performed all experiments. All authors analysed and discussed the data. J.L., F.Y. and Z.C. wrote the paper.

