## SUPPLEMENTARY INFORMATION for "Clostridium ablates tumors by degrading collagen mesh and inducing ferroptosis"

**Table S1. Collagenase associated proteases of RJ-1.**

| Sequence _ID | Gene_Sequence | Annotation | Gene Name |
|---|---|---|---|
| PROKKA_00944 | ATGCAAAGAATAGAATTATTAGCACCAGCAGGTGATTTAGAAAGATTAAAAACAGCATTTATA<br>TATGGAGCGGATGCCGTTTATATAGGTGGTGAAATATTTGGAATGAGATCTGCTGCTAAGAAC<br>TTCAATAAAGAAGATATGGCAGAAGGTGTTAAATTTGCGCATGAAAGAGGCAATCAAGTATT<br>TGTTACTGTAAATATAATACCTAGGAATGAAGAGTTTGAACAGTTAGAAGCTTACTTAAAAGA<br>ACTTGAAGAAATAGGTGTAGATGCAGTTATAGTAAGTGACCCAGGAGTATTTAGCGTAGTAA<br>AAAAAGTTGTCCCAAATATGGAAATACATATAAGTACTCAAGCTAGCACAACTAATGCTGCAT<br>CAGCTACATTTTGGTATAATCAAGGAGCAAAAAGGGTTGTTATGGCAAGAGAGTTATCTTTT<br>GAAGAAATAAAAGAGATAAGAGACAACTCTCCAGAAGGTATGGATATAGAAGCATTTATACA<br>TGGCGCTATGTGCATGTCTTATTCAGGAAAATGTGTAATAAGTAACTATACAACAGGTAGAGA<br>TGCTAATAGAGGAGCTTGTGCACAATCTTGTAGATGGAAGTACACATTGGTTGAGGAACAAG<br>AAAATGGAGACTATGAAAAAGTTTTAGATGACGTAGATGCAGAATTTTTCTTTAATACAAAA<br>GATATGTGTATGATAAACTATATTCCACAAATTATAGAAAGTGGAATAAATAGTTTTAAAATAG<br>AGGGAAGAATGAAAACTGCATATTATGTAGCAACTACTGTTAGGGCTTATAGAATGGCTATAG<br>ATGAGTATATAAAAAATCCTGAAAATTGGAAGTTTAATCCGATGTGGTTAGAAGAACTTAAA<br>AAAGGAAGCCATAGACATTTTAGTGAAGGATTCTATTTAGGGAAAACATCTACTAGAGATCA<br>AAACTATGAATCTGCATCGTATGTAAGAAACTATGATTTTATAGGTGTAGTTAGAGGACATGA<br>AGAAGAAAGTGGACTAGTTATAGTTGAACAAAGAAATAGAATGTTTGTAGGAGATGAAATA<br>GAAATAATAGGACCATATAAAGAAACTATGTATGGCAAAATTTTAGAAATGTACAATGAAGA<br>AAATGAGCCGATTGAATCTGCTCCACATGCAAAACAAATAGTAAAAATGAAATTAGACATAC<br>CGGTTGAAGAACATTATATGTTAAGAAAGCCAATAACAACTATAAATGTATTATAA | CDD:223896 COG0826, COG0826, Collagenase and related proteases [Posttranslational modification, protein turnover, chaperones]. | - |
| PROKKA_01922 | ATGCATAAAGTTGAGCTTTTATCATCAGCTAAAGATTTAAATAGCTTGAAATTAGCGTTTGAA<br>AGTGGTGCAGACGCTGTTTATATAGGTGGAGATTCATTTGGAATTGATGCTATAGCTAAAAAC<br>TTCTCAAGAGAAGAATTACAGCAAGGCGTTGAATTTGCCCATCACAACAATAAAAAAGTTTA<br>TGTAGCTGTTAATGTTATGCCACACAATGAGGACTTCGATAGTTTAAAGGATTACTTAGTAGA<br>ACTTGAAAAGTTAAACATAGATGGAATAATAGTAACAGATCCTGGGGTTTTAACAATAGCTA<br>AAAATACTATACCTAATGTTAAAATTCACATGGGACAGCAAGCAAACGTAACTAATTTTAATT<br>CAGCTAACTTTTGGTATGACCAAGGAGTTAGAAGGATAATTATCACTAGTGAATTATCTTTTG<br>ATGAAATAAGTACAATAAGAGCTAAAACTCCATTAGATATGGAAATAGAAACTTTTGTCCATG<br>GACCTATATGTATATCTTATTCAGGAAGAAGGCTTTTAAGTAGCTACTTAGGAAGTAAAGATA<br>ATGTAGAGTACAAAGAAGGAAACAATAAGTATAACTTATTAGAAGAAAAAAGACAAGGGGA<br>ATACTACCCAGTTTTTGAAGATGAAAAAGGAACTTTCTTATTTAACTCAAAAGACCTATGTAT<br>GTTAAACTTTGTACCTGACTTTGTAAAAGCAGGTGTGACTAGTTTAAAGATAGATTCAAGATT<br>ACAAAGCGACGAATACTTAGAAACAGTTATAAAAGTCTATAGAAAAGCTCTAGATGAATTTT<br>ATAAAAACCCTAATGAATGGGAATTCAATCCTATTTGGTTAGAGCAGATAAAAGAAAAAAGT<br>CATAGACCATTGACACATGGATTCTACACAGGCGAAGCTATGCCAGATGATTATGAATAA | CDD:223896 COG0826, COG0826, Collagenase and related proteases [Posttranslational modification, protein turnover, chaperones]. | - |
| PROKKA_02733 | ATGAATACAGAATTACTAGCACCTGTAGGTAGTTTTGATGCGTTAAAAGCAGCTGTACAAAAT<br>GGAGCAAATGCAGTTTATTTAGGTGGAAAAGAATTTAGTGCTAGAGCCTCAGCAAATAACTT<br>TGATAGAGAAGAATTAAAAGAAGCTGTAAAGTATGCGCATATAAGAAATGTAAAAGTATTTG<br>TAACGGCAAATACGCTTATAAAACAATATGAAATAGATGATTTTCTTAATTATATAAAATATTTA<br>TATGATATAGATGTTGATGCAGTTATACTTCAAGATATAGGAATGGCTAAACTAATAAAAAGTT<br>TACTACCAGACTTTGAATTACATGCAAGTACACAAATGGTTGCTCACTCTCTAGAAGATACTT<br>TGTATTTACAAAACATTGGATTCGATAGAGTTGTTTTAGCTAGAGAACTTAATATAGATGAAAT<br>TGAACATATATGTAAAAATACTGATGTTGATATTGAAATATTTGTACATGGAGCATTATGTGTAT<br>GCTATTCAGGTCAGTGCTTAATGAGCAGTATGATAGGAAATAGATCTGGAAACCGAGGTAGA<br>TGCGCTCAACCATGTAGACAAAAGTATGAATTAATAGACATAAATTCAGGTAAAGTTATAGAT<br>ACAGAAGGCGATTATTTATTAAGCCCTAGGGATTTAAATACAATAGAGGAAATAGATAGAGTT<br>ATAGAAGCTGGAGTACACTCTTTAAAGATTGAAGGAAGAATGAAACGTCCAGAATATGTAGC<br>AACTGTAATAAGTAGTTATAGAAAAACTATAGATGCATATTTAGAAAAAAATCAAGTAAATGT<br>ATCAAATGAAACTATGAACAATTTATATACTATATTCAATAGAAAGTTCACAAAAGGATATTTA<br>TTAGGAGAGGTTGGAGAAGATGTTATGAACTCAAATCGACCTAATAATGTAGGACTTTATATT<br>GGAAAAGTAATTGATTACAATAAAAAAGCTAAACGCTTAAAAATTAAATTAGAGAATACACT<br>TAAAAAAGGTGATGGAATTAACTTAGGTGGAGGAACTATAGGAAGAATTATTAAGCAAAATA<br>ATACAATAAGTGATATAGGAGAAGCTGGAGAACTTATAGAATTAGATTTTATAGGAGAAGCTA<br>AAAAAGGACAAATCGTATATAAAACATCAGATAGTGATTTATTAAATACTATTAAGAAATCATT<br>TGAAGAAGATGTTGAAAATATAAAAATTCCTATAGATGCGACTATTGAATTAAAGCTTGGACA<br>ACCTCCAAAACTTAGACTAAAAGATGGGTATGAAAATGATATAACTGTTACAAATGAAAAGC<br>TTGTAGAGAAAGCTATGAAGGTTGCTCTAGGTGAGGAAAAAATAGTATCTCAAATAAAGAAA<br>CTAGGAAACACAGCATATGTATTGAGAAATATAGAGGTTGATATAGATGAAGATATAAGTTTA<br>CCAATAAGTATATTAAATCAACTTAGAAGAGATGCTATAGATAAATTAAGTAATGAAAGAGTC<br>ATTATAAAAAAATAGAAACTTCAAAGATTCTTTTATAAAGTACTCACCAAAAGTTATAAAAGAG<br>GATAAAGACATAAAACTAAGAGTTAAAGTTAAGAATATAGAACAATTAAAAACTGTATTATTA<br>TATGATATAGATGCAGTATATTATGAAGACATAAACACATTAAAAGAAGCTAAAGAAGCTTGT<br>TACAATAATATAAAACTTATCTACTCTTCTCCTAGAATTCTTAGAAATAAAGATTATGCTATTTT<br>AAATAAGGTTGAAAAAGTACCTGTACAAGCTGGAAATTTAGGTTGTGTAAATTTATATAAGG<br>GAAAAGATATATATATAGATAGCTACTTAAATTCTTTTAATAGTGAAACTATAAATCACTATAAA<br>TCAGAAGGCGCAAATACGATTTGTTTATCTCAAGAGTTAAATTTAACAGAAATAAAAGATATG<br>TTAAAGTACACTGATAGTGAAATAGAAAGCATAGTTTATGGATATACACCTCTTATGATTAGTG<br>AATACTGTCCAATGGGAGTTTTAGTTAGAAACTGTAAAAAAGATAAAAGAAGTTCTATTTGC<br>AACAAGAGTTTATATGCACTAAAAGATGCTAAAGATGGGACTTATAGATTATCTCAAGATATT<br>TTCTGTAGAACTAGTATATACAATTCTAAACCACTATGTGTATTAGAAGATTTAAGTGAATTAA<br>ATAAAGCTGGCATAAATGTATTTAGAATAGATTTAACATTTGAATCAAATGAAGAAATAAAGT<br>CTATAATTGAAGCTTTTGAAGAGTGTATAGAAAATGATTTTAATATAGGCATAAAATCTAAAA<br>AATTATATAAAGAGCTTGAAAATACTGGATTAACATCCGGTCATTACTATAAAGGTGTTGAATA<br>A | CDD:223896 COG0826, COG0826, Collagenase and related proteases [Posttranslational modification, protein turnover, chaperones]. | - |
| PROKKA_02921 | ATGTTTAAAAGAAATTTAGCTATTTTAACAATTGCAGGTTTATCATGCGGATTTTTAGCACCTA<br>CAGTAAGTTTTGCAGATAGTCAAACTAAGGAAGAATTGCCTCAAGCTTATAATGTTACAGCA<br>GATAAAGATTTAAAAGATGTATTAAAAAATGTATCTGAAGATTACAAAGTAAAAGAAAATGC<br>AGATCAATATGAATTAGTTAAAAAAGAAAAAGATGACTTAGGATTTACTCACTATACTCTAAA | CDD:225768 COG3227, LasB, | Gelatinase |

| | | | |
|---|---|---|---|
| | ACCTAAAGCTGATGGTTATTTTGCAGATAAAGCTGAGGTTAAGATACACACAGATAAAGATG<br>GAAAAGTTGTTTTTGTAAATGGGGATTTAGATCAAGGTAAATTGGAAGTTAAAAATGAAACT<br>AAAATAGATAAAGATAAAGCTATTGAGTTAGCTTTTAAATCAATAGAAAAATCACGTGATGAA<br>GTTAAAAACTTATCAGGTGAAGATACAATTCAAGATGCTAAAATTGTTGTTGATGAAAAAAC<br>TAATAGAGCTGTATATGCTTTAGATTTATCATATTCTGTTCCAGAGGCTGCTCATTGGTTAATTA<br>AAGTAGATGCAGAAAATGGTAGTATAGTTGAAAAACAAAATGTTTTAGAAGAAGCAATTCA<br>AGCGACAGGTACTGGAATTGGATCAGATGGACAAGTTAAAAACTTAAATATAACTGAAGATA<br>ATGGTAAATATCAATTAGTGGATACTACTCATAAAGGTCAAATAATAACTAAAAGTTTCGAAG<br>CTTTTAAAGGTGATACAACAGTTGGAAATATTATTACAAATATAAAAAACTTCTTCGATCAAG<br>ATAAAGCAGCTGTAGATGCTCACTATTTTACAAATCAAGTTTATAACTACTACAAAGATACTC<br>ATAATAGAGAAAGTTATGATGGGAATGGCGCAGATATAAATTCTTATGTTCATGTTCCTGAAG<br>AGGATGGAAGTAGTATGAGTAATGCTTACTGGAATGGCGTAGAGATGAGTTATGGTGATGGA<br>AATAAAACAACAGAAAATGCTTTTACAGCAGCAAATGACGTTGTTGCACATGAGATAACAC<br>ATGGGGTTACATCATCATCAGCAGACTTAGTTTACCAATATCAACCAGGAGCATTAAATGAAT<br>CATTCTCAGATGTATTTGGTTATTTTGTAGATAGTGATGACTGGCAAATGGGAGAAGATTTATA<br>TAAAACACCAGGTGAAGCTATAAGAGATTTACAAGATCCTACAAAACATGGACAACCAGCA<br>AATATGAAAGACTATAAAAACTATAGTATATATTATGATAGAGGTGGAGTTCATATAAATAGTG<br>GTATACCTAATAAAGCAGCATATAACACTATAACTAAAATAGGGAAAGAAAAAGCAGAGAAA<br>ATATATTATAGAGCTCTTACTCAATACTTAACAAGACAATCACAATTTAAAGATGCTAAAAGTT<br>CTTTAATTCAATCAGCAAAAGACCTTTATGGTGATCAAGTAGCTAATGATGTTAAAGCTGCAT<br>GGGATCAAGTTGGAGTAAAATAA | Zinc metalloprotease (elastase) [Amino acid transport and metabolism]. | |
| PROKKA_03239 | ATGAAGAAGAGACATTCTAAGTTTTTGATTTTTACTTTAGCTTTAGCCTTGGTTTTAACTAATT<br>TAACATTCATTTCATTTGCAAATACTCAACCACCTAGTTCTAGTAATATTCCAGGTGATATAAA<br>TACTATGCCTTTCCATATAGGTCAAAAGATTAAGCACGATACTAAGGAAAATCAGTTCAAGAA<br>AGATTTAAATATTGAGCCTTTAGGAAAGCAAGATACAGACACAAAACTTAAGTCAGGGAAG<br>TCCGCAAAGCTAAGTAAGTCTGCTACCTATACCTATAATGACCTTAATAAATTAAGCTATAAGG<br>AGTTAACAGATTTACTAGTTACGATAGATTGGTTTCAAATAAAAGATATTTTTGTTTATAATGA<br>AGATGCTTACAAATTTTATAACGATGAAGCTCGTGTTAATGCAATTATTAAGACGTTAGAAGA<br>TAGAGGAGCAACTTATACTGCTACTGATGATAAAGGAATACCTACTTTGGTAGAAGTTTTAAG<br>ATCTGGTTTTTATTTAGCTTATTATAACGATTCACTTAGTTATTTATACGATAGAGCTTATAGAG<br>AAAAATGTATACCTGCCATGATTTCAATTCAGAAAAATCCTAATTTCAAGATTGGGGAGTCTG<br>GTCAAAATAATGTTATTCAGTCTTTGGGTATGTTAATTGGAAATAGTGCTTGCAATGTTGAGG<br>TTATAAATAATTGCGTGCCTATTTTAAATCAGTTTAATAGCAGTTTAGATGCTAATTTAAAAGA<br>TTACTCAAAAACAAATGCTGTATTCCAAATCACTAAGGGTATAGAATATGATTTAAATTCCTAT<br>TTGTACAATTCTAATACAACTGCAGACAAATCACCTTGGTTTTCTAAAGTTGATGGATATATAG<br>ATGCTGTTACAAACTTTGCTATATTAAGCAATGTAACTGATGATAATGATTGGTTAATTAATGC<br>AGGGATGTATTATTGCGCTAAATTATCGAACTTCCATTCAAATTCAACAAATATTCAAAAGAA<br>ACTTGATAGCTGTTTAGAAATTTATCCATATTTATCTATTCAATATTTTGATGCAGTCGATTTTAT<br>TTCATATTACTTCAATGGGAAGTTAGTTGATGGAACTGTTCTAGATATTAATAAAATAAGAGA<br>AGAGGGAAAATCACACTATTTACCTAATGTGTATACTTTTGAAGATTCATCTATGGTAATTCGT<br>ACTGGTGATAAGGTTACGAAGGAAAAAGTTCAAAGATTATATTGGGCATCTAAAGAAGTAAA<br>AGCTCAATTCCATAGAGTTATAGGGAATGATAATGAACTAGAAAAAGGAAATCCAGACGATG<br>TTTTAACTATGGTTTTATATAATAGCCCTAAGGAATATAAACTTAACCAACCTCTATACGGATAT<br>AGCACTGATAATGGTGGTATGTATATAGAAGGTGACGGCACATTTTTTACTTATGAGAGAACT<br>CCTGAGGAGAGTATATTTAGTTTAGAAGAGTTATTTAGACATGAGTACTGTCACTATTTACAA<br>GGTCGTTACATGGTTCCTGGTATGTGGGGAACGGGTGATTTTTATCAGGGTAAGGATTGGAG<br>ATTGACTTGGTTTGAAGAAGGTAGTGCAGAATTTTTTGCAGGGTCTACTAGAGATAATGATGT<br>ACTTCCTAGAAAATCTCAAGTTTCTGGTTTAGCAACTGACCCAGCGGAAAGATTTTCTACAG<br>ATAAATTACTTCATTCTAAGTATGGATCTTGGGATTTCTATTATTATGGTTTTGCATTTTGTGATT<br>ATATGTACAATAATAGATTAGATATATTTAATAATTTAGTTGATAACATTATATCTAATAATGTATC<br>TGGATATGATTCTTATATAGAAACATTAAGTAAAAGTACTACAGTAGATCAAGGTTATCAAAC<br>TCATATGCAAAACTTAGTTGATAAGTATGATTCTTTGACAGTTCCACTTGTTTCTGATGATTAT<br>TTGGCTACTCATGATGCTATTGATTTAAATAAAATTTCTTCTGATATTAAAAATATAGTTCCATT<br>AAAAAATGTAAATATAGTTAGAGAAAAAGGCGAATTTTTTGATACATTTACTTTAACAGGAAC<br>TTATACAGGAGATACTTCTAGAGGAGAAGCTAAAGATTGGGAAGCTATGAATAACAAATCTA<br>ATGAATTTTTAAATTCCTTAGAAAAACTTCCTTGGTCTGGTTATAAGACTGTTACTAATTACTT<br>TGTTGACTACAAGGTTAATAATAATAAATATGAATTTGAAGTTGTGTTTACCGGTATTTTACCA<br>AAAGGTGAAGTTGTAGAAGGTGATATTAAAGAAATTGAGCCAAATGATAGTTTTGAAACTGC<br>TAATACAATATCTTTAGGAAGTGAAGTAAAAGGTTCTTTAAGCAAAGATGATACTAATGACGT<br>TTTTTCTTTTGAAATTAAAGATCCTAAAACTATAGATATAGTTTTAGATAATTTAGATGGTCATG<br>GTATTAACTGGATACTATATAAGTCTACTGATTTAAATAATTATGTTGCTTACCCTAGCATTGAA<br>AACAATTTATTAAAAAATACTTATAATGCAGATGCTGGTTTATATTATTTATATGTCTATAGTTAT<br>GACTCTTTAGATAGTAACTATACAATTAATATCAAATAA | CDD:203949 pfam08453, Peptidase_M9_N, Peptidase family M9 N-terminal. This domain is found in microbial collagenase metalloproteases to the N-terminus of pfam01752. | *Kappa* toxin |

**Table S2. DNases of RJ-1.**

| Sequence _ID | Gene_Sequence | Annotation |
|---|---|---|
| PROKKA_00007 | ATGAGTTCTCCTAAGTGGACAGACGAACAACAAGCTGTAATAGATAGCAGAAATTGTAATCTTCTAGTAGCTGCTG<br>CTGCAGGTTCAGGGAAAACAGCAGTACTAGTTGAACGTATTATACAGATAATAACAGACACTAAAAAGCCAGTGG<br>ATATAGATAAATTATTAGTAGTTACATTTACAAATGCAGCTGCTTCAGAAATGAGAGAGCGTATAGGAGATGCAATAG<br>CTAAGGCATTAGATAAAAATCCTGAAAATAGCCATTTACAAAATCAATTAGTTTTATTAAATAGAGCAAGCATTACA<br>ACTATACATTCATTTTGTTTAGATGTAATAAAATCAAATTTTCATAAAATAAATATAGATCCTAACTTTAGAATAGGAG<br>ATCAAACAGAATGTTCTCTTTTAAAGCAAGAATGTATAGAAGAAATTTTTGAACAATACTATGAATCAAGAGATAAG<br>GGATTTTTAAATTTAGTGGAAAGCTATGCAGAAAAAAGAGGAGATAAAGATTTACAAGATATAATATTATCTATATAT<br>AACTTCTCGATGGCATCACCATACCCTAAAAAATGGTTAGAAGATTCAGCTGAATTATTTAATATAAATGATGATTTT<br>GATTTTTCAAATTCAATATGGTCAGAATCAATATTAAGTAATGTAAAAATAGAGATAGATGGAATAGCATCTAGTATG<br>AAAAGTGCGATTGAATATGTAGATGGAATTGACGAATTAGAAACTTATAAGGAAAAATTAAATATAGAGTATTCTCA<br>AATATTAAATATATTAAGTGCTTGCAATGAAGGCTGGGATAAGACATATTACTTTATGAGCAATATGCAATTTGAAAA<br>CTTCGCAAAAGGAGTTAAGAGATTAGGAAAAGATACCCCTGACTACATAAAAAAAGCAAGAGAAAATGCTAAATC<br>TATAAGAGATAAAAATAAAAAATCATTAGAGAGTATAATAGGATCAACTTTTTATAAAAGTAATGAACAGATTTGTC<br>ATGAAATAAAGTATTTATATCCAGTTGTAACTTCTATTTCAAACCTTATAATAGCGTTTGAAGAAAAGTATCAAGAAA<br>AGAAAAGAGAAAAAGGAATTATAGATTTTAATGATATAGAGCATTTTGCATTAGCTATATTAACAGATGAAGATGAA<br>GATGGAAATATAATACCATCAGATGTAGCTAAGATATATGTAGAAAAATTCTCAGAGATATTTATAGATGAATATCAA<br>GATAGTAACTTGGTTCAAGAAGTTTTATTAAGTACTATCGCTAGTATTGAAAATCCAAATCGATTTATGGTTGGAGAT<br>GTAAAACAAAGTATATACAGATTTAGACAAGCAAAACCAGAAATATTTTTAGAAAAATATGCAACTTACGATACAG<br>AAGAAGGTACTAAAAATAGAAAGATAATGCTATATAAGAACTTTAGAAGTAGGAAAGAAGTTGTAGATTGTGCAAA<br>CTATATTTTTGAAAATATAATGAGTAAAAATATAGGTGAAATCGAATATAGCGAAAAAGAAAGATTAAATTTAGGCG<br>CAAGCTTTAAAGAGTGTGAAGAAGAAAATGTGTTAGTGGGAGGACCTGCCCAAATACATTTAATACAAAAAAATAT<br>TAAAGCTAAAAATGAATCAGATGAATCTGAGGAGAATGAGGAAGAACAAGAAGAAATAGATAACATACAATTAGA<br>AGCTAGAATGTTAGGAAATATAATCAAAAACTTAATGAAACCCAATGAAGATGGGAAGGTAACAAAAGTTTATGAT<br>AAAAAAATTGATGGATATAGAAATGTAGAATTTAAAGATATAGTAATACTATTGAGAGCAACATCTAGTTGGGCACC<br>TGTATTTGTTGAAGAGCTAATGAATATGGATATACCTACATATGCAGATACAGGTATTGGATATTTTGATACTATAGAG<br>ATAAAAACAATACTAGCATTACTTCAAATAATTGATAATCCAATGCAAGATATTCCTTTAATTGCAGTATTAAAGTCA<br>CCTATGTTTGGGTTTACACCAGAAGAGCTAATTGATATAAGAATAGAAGATAGACAGAAAAGTTTTTATGAAGCTTT<br>ACAAATAGCAGGTAGTAAAGATGGAGATTTAGGTGAAAAAATAAAATATTTCTTAGATAGATTAGATAATTTTAAAA<br>ATAAATCACTTTATATGAGTACTGATGAATTTTTATGGTACATATACATAGAAACTGGTTATTATGCATATGTAGGAGC<br>ACTACCAGCAGGAACACAAAGACAAGCTAATTTAAAGGTATTATTTGAAAGAGCTAAGCAATTTGAATCAACAAGC<br>TTTAAAGGGATTTTCAATTTTATAAACTTTGTAAAGAAATTAAAGAGATCGAACTCTGATATGGGAAGTGCTAAAAC<br>ATTAGGAGAAAATGCAAATGTAGTAAGAATCATGAGTATACATAAGAGCAAAGGGCTAGAGTTTCCTATAGTTATAT<br>GTGCAGGAATGGGTAAAAATTTCAATACTCAAGATTTTAGAAAAGACATTCTATATCACCATAGGTTAGGGTTTGGC<br>CCTCAGCTAGTAGATTATGAGAGAAGAATATCATATCCTAGTATAGCGAAAGAAGCATTAAAAACAACTATAAATAT<br>GGAAAACTTATCAGAAGAAATGAGAGTTTTATATGTTGCATTTACACGACCTAAAGAGAAGTTAATCATAACAGGTT<br>CGGTAAGAGATATAGAGAGCAGCATAAAAAAATGGTCTGAAGATCTAGATAATGAGGGACCTGTTTCAGAATATCA<br>AATTTTAAAAGGAAAGAACTTTTTAGATTGGATAATGCCTGCAGTTATAAAGCATAAAGACTTAGATGAACTACGCA<br>GTATAATTGAATTAGACGCTAATTACTTAAATAATCATGAATCAAATTGGGAAACTAGAATATGGGAAAGATCGGATA<br>TATTAATAGAAAAGACTTCTAATGATGAAGAGGAAAGCATAGATACAATTTTATCTAATTTAGATATGAGCGAAAATG<br>AAAGTGAATTTTATGAAGATATTAAATATAAGCTAGATTATAAATATCCATTCATGGAATCTGTTAATAGGGCAGGAA<br>CTATATCTGTTACTGAAATAAAACGATTAGAAAATAAAGCAGAGATTGATTATAGTGCTCAAGAATTAATTGAAAAG<br>CAAACTGAAATAAAAACTCCGTTATTTATACAAGAAGATGTAAGTAAAAATAAATTAACTGGAGCTCAAAAAGGTA<br>CTATAATGCACTTATTCATGCAAATTGTTGATTTAACTAAAGTGAGCAATTTAAATGAAATTAATGAAGAAATTAATT<br>TAGCTGTAAAAAAGAAAGTATTAACACAAACTCAGGCAGATACTATAAATCCATATAAAGTATTAAAGTTTTTTAAA<br>TCTAATTTAGGAAAAAGAATGTTAAGTTCTTATTATTTAAAAAGAGAACAAGCTTTCTATTTACAAATAAACATGAG<br>TGATATATTCAAAAATGATGATAGAGTAGAAATTATAGAAAATGAAGATAAACTTATGGTAAGAGGGATAATAGATG<br>CTTACTTTGAAGAAAATGATGAACTTGTTATAGTAGATTATAAAACTGACTTTGTAAATGAAGAAAATAAAGATACT<br>ATTACAGAAAAATATACTAAACAGCTTGAAATATACAGTAGGGCGTTGGGTACATTAACAGGCAAGAAAGTTAAAG<br>AAGCATATATATACTTATTTGGAACTGAGGAAGAAGTTTTATATAAGTTTGATTAA | CDD:224000 COG1074, RecB, ATP-dependent exoDNAse (exonuclease V) beta subunit (contains helicase and exonuclease domains) [DNA replication, recombination, and repair]. |
| PROKKA_01654 | ATGAAAATTATAACTTACAACATTCATAAAGGTATGGATGCACACAATAAAGATACCTTAAATGAATTATTAAGTTTT<br>TTAAAAAAGATAAATGCAGATGTAATATGCCTTCAAGAAGTTTTAGAGAGTATACATTATAAAATTTTAAATGATTTA<br>AATATAAAAGGTTACTTTTTTAAAACTGTAGAGTTAAAAGGTGATAATTATGGAATTGCTATATATTTTGTTGAATGTG<br>TAAAGTACACAGATGGATTTTATTTAAGTAGTAAAAAAGAGCAAAGGGGATTTATTCATACTCAATTTTTTATAAATA<br>ACAAATTAGTTAATATAATAAATACTCATTTAGGGTTAAATGAGGAAGAAAGAAAAATACAGATTGAGGAAATTTAT<br>AGTTATGTAAATAATTTAAGAGGAAAAATTATTATATGTGGTGATTTTAACCAAACGAATATAAAAAATAAAAGATTAT<br>TATGACTTATCTATATTGTTTAGTTGTGAAAATATAGAAACATTTCAACCTAGTAAATCTAGAATAGATTATATATTTAT<br>GTCTAAAAATCTAAGACCGAGTAGATATAAAGTTGAATTTATTAATTTATCTGATCATTATCCTGTAATAGGAGAATTT<br>ATTATATAA | CDD:197306 cd08372, EEP, Exonuclease-Endonuclease-Phosphatase (EEP) domain superfamily. |
| PROKKA_03275 | ATGGAAAAGATTGAAGGAATGGTAAGTGATATTATATTTAAAAATGAAGATAATGGATATGTTATAGCTCATTTATCA<br>AATGAAAATAACGATATAGTTATAGTTGGATGTATGCCTACTTTAACAGTTGGAGAGAGTATAGAAGTTGAAGGTAA<br>GTGGATAAATCATAAAACTTATGGAACTCAATTTGAGGTTAACTCATTTCTGCCAGTGACACCATCATCTATAGAAG<br>GTATATATGTATACTTATCCTCTGGAGTAATAAAAGGGATAGGAGAAAAGATGGCAAAGAGGATAATAGAAAAGTTT<br>GGAGTAGATACCTTGGATATTATTCAAAATTCTCCACATAGGCTTTCAGAAGTTGAGGGTATAGGGAGCAAAAAAAT<br>TGAGCAGATAGCTAAAAGTTATGAAGAAAATAGAGAATTAAGAAATATAATAATGCATTTATCTCCTTATGGAATAAC<br>TCCTAATTTTTGTTTAAAGATATATAAAAGATATAAGAATAAAGCAATAGATATAATAACTAAAAACCCATATAAGTTA<br>GCAGAAGATATAAGAGGAGTAGGATTTAAAGTAGCAGATAAAATAGCGTCAAATTTAGGAATAGATAAATTTTCTAA<br>AGATAGAATAATGCAAGGTATCTTATACACATTAAATCAATCTTTAGGAAGTGGTCATACATATTTACCCAAGGATAT<br>CTTAATAGATGAAGCTAGCAAGTTGTTAGGGGTAGATAATGAATATATTTCAGATTCAATTTTAAGCTTAGCTTATGAT<br>CAAAAAATTCATATTGAAAGAAATTTTGGAGATGAAGAGCACATATATCTAATACCATATTATCTATCTGAGAATGGA<br>GTTTGTAAGCAAATCATAAAATTATCTCAATCTCAATTTAAAGAATTAAATATAGATATAGATGAAGAAATAGAAAAA<br>ATAGAAAAAGAAACTGATATAAAGTTAGCCACAAATCAAGTATTAGCAGTAAAAGAAGCTGTTGAAGGTGGATTAG<br>TAGTTATTACTGGAGGTCCCGGAACAGGAAAGACGACGACTATAAATACTATAATAAAAGTATTTGAAAATAATAAA<br>CAAGAAGTTCTTTTAGCCGCACCTACAGGGAGAGCAGCAAAGAGGATGAGTGAGACTTCTAATAAAGATGCGAAA<br>ACAATTCATAGGCTTTTAGAAATGGGATATGCAACAGATTCAGAAGAACTGGTTTTTTTAAAAGATGAAGAAGACC<br>CTATAAAAGCAGATGTAGTTATTGTTGATGAAGTTTCAATGGTAGATATTGTTTTAATGTATAGCTTATTAAAAGCAAT<br>AAAACCAGGAACTAGATTAATTTTAGTTGGAGATAGTGATCAGTTGCCATCTGTTGGAGCTGGAAATGTTTTAAAA | CDD:223581 COG0507, RecD, ATP-dependent exoDNAse (exonuclease V), alpha subunit - helicase superfamily I member [DNA |

| | | |
|---|---|---|
| | GACATTATTGATTCAGATGTTATAAATGTTGTTAGATTGAATGAGATTTTTAGACAAGCTCAGGAAAGTATGATAGTT<br>GTAAATGCACATAGAATAAACAAAGGATTGCCACTTCACCTAAATGTTAAAGGGAAAGATTTTTTCTTTATAAAAAA<br>AGATACAAATGAAGCAATTTTAGAAGAAATAGTTGGCTTAGTTAGCCAAAGGCTACCTAAATTTTATAACTTAGACA<br>AGCTACAAGATATACAAGTTCTTGCACCTATGAGAAAGGGTGATTTAGGTGTTACAAATTTAAATATAGAGTTACAA<br>AAATATTTAAATAAAGAAGAAAAGTTTAAAGTTGAAGAAACACTACAAAAAAGGATTTTTAGAGTTGGCGATAAA<br>GTTATGCAAATAAAAAATAATTATACTAAAAAATGGGAAAATGAAGATAAGAGTGAAAGTGGAGAAGGTATATACA<br>ATGGAGATATAGGTTATATTTATCATATAGATAAGGAAAACAAAATTATATATGTTCTATTTGATAAAACTAAAATAGT<br>GTCTTATGAATACTCTGAGTTAGATGAACTGGATCACAGTTTTTGTACGACAATACACAAAAGTCAGGGAAGTGAA<br>TTTCCAGCTATTGTAATTCCAGTAACTTGGGCTCCGCCTATGCTATTAAATAGAAATCTACTTTACACAGCAGTAACT<br>AGAGCTAAACAATTAGTAGTTTTAGTTGGAGATGTTAAATACTTAGAATTTATGATAAAAAACAATAGGATAAATGAT<br>AGACACTCTAATTTAAGTAAAAAATTGAATAGATTAAAAAAAGAAGGGGTGCTGATAAAAAATTAA | replication, recombination, and repair]. |
| PROKKA_03499 | ATGTTATTTGACTCACATGCACATTTAAATGATGAAAGATTTGATGAAGATAGAGAAGAATTAATAAACTCACTTAA<br>AAATAAAGGTGTAGATTTAGTTTTAAATCCAGGCGCTTGTATAGAAACATCTAAAAGCAGTGTAGAATTAGCTAATA<br>AATATGATTTTATATATGCTGCAGTTGGAGTTCATCCTCATGATGTAGGTGAAATGAACGAAGATGATATAGAAACAT<br>TAAGAAAACTTGCTACTCAAAATGAAAAAGTAAAAGCTATAGGTGAGATTGGTTTAGATTACTATTATGATAACTCT<br>CCAAGAGAAACTCAAAAGGAATGGTTTAAAAGACAAATAGAACTAGCTAATGAACTTAAATTGCCTATAATAATTC<br>ATGATAGAGATGCACATGGAGATACATTTGAAATAATAAAAAACACAAAAAGTCCTGAAATAGGATGTGTTCTTCAT<br>TGTTATAGCGGGAATGTAGAACTTGCTAAAGAATATATTAAAATGGGATGTTTAATATCTATACCAGGAACTGTAACA<br>TTTAAAAATAACAAAAAAACTAGAGAAGTTGTTAGAGAAATACCTCTTGAGTATTTACTTATAGAGACTGATTCTCC<br>ATATATGGCTCCAGAACCACATAGAGGAAAAAGAAATGATCCTTCTTTAGTTCAATTTGTAGCTGATAAAATAGCTC<br>AAGAAAAAGGAATTTCATATGAGCAGGTTTGTGAAGTTACAAAAGAAAATGCTAAAAGATTTTTCAATATAAAATA<br>A | CDD:223162 COG0084, TatD, Mg-dependent DNase [DNA replication, recombination, and repair]. |

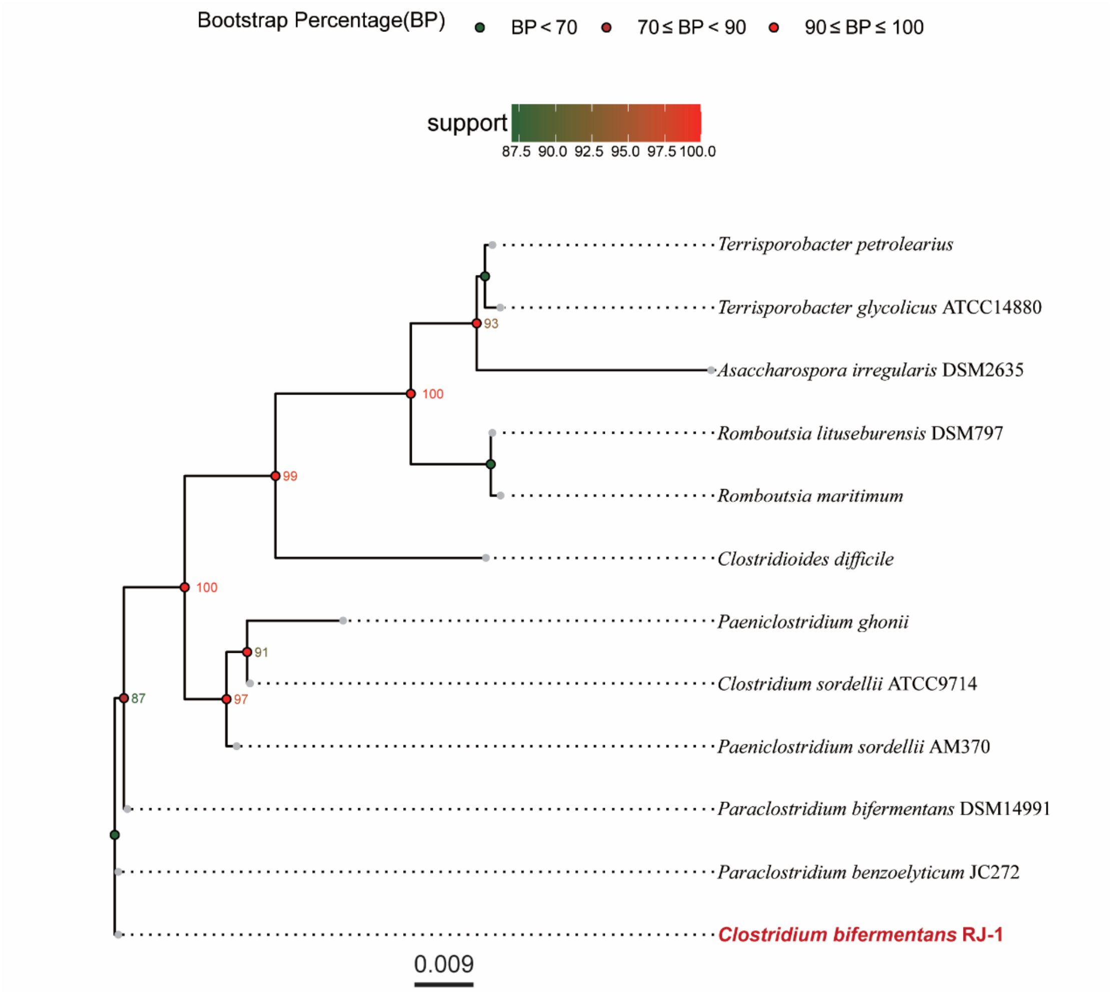


**Fig. S1 Phylogenetic tree based on the data from 16S ribosomal RNA sequencing.**

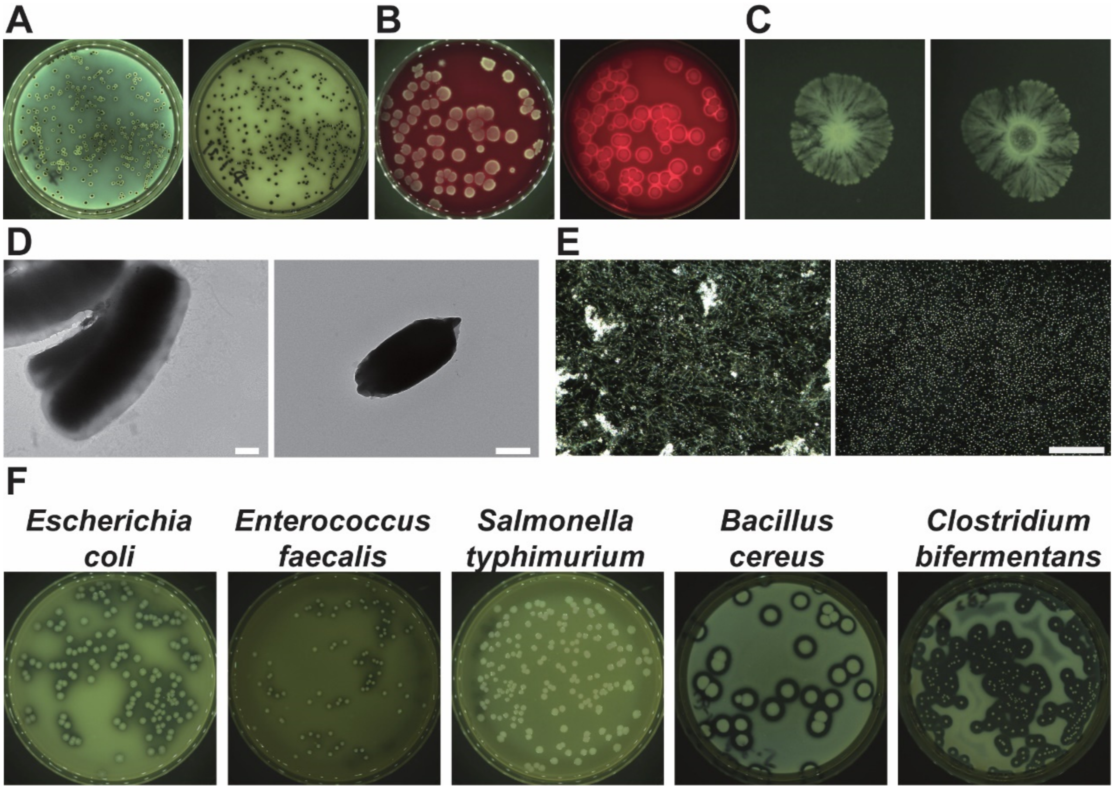


**Fig. S2 Microbiological characterization of *Clostridium bifermentans* RJ-1.**
(A) RJ-1 colony morphology on differential clostridial agar. Left, no background color. Right, white background. (B) RJ-1 colony morphology and hemolysis on Columbia blood agar plate. Left, incident light photography. Right, transmitted light photography. (C) Dynamic colony morphology changes of RJ-1 on BHI plates with 1% agar. Typical branched colony forms by bacterial swarming motility at day 7 (left), and obvious autolysis in the center of the colony appears at day 10 (right). (D) Transmission electron microscopy (TEM) pictures of RJ-1 vegetative cells (left) and spore (right) (Scale bar: 500 nm). (E) RJ-1 vegetative cells (left) and spores (right) morphology under dark field microscopy (Scale bar: 100 μm). (F) Skim milk plate assays for collagenase activity of different bacteria strains as indicated.

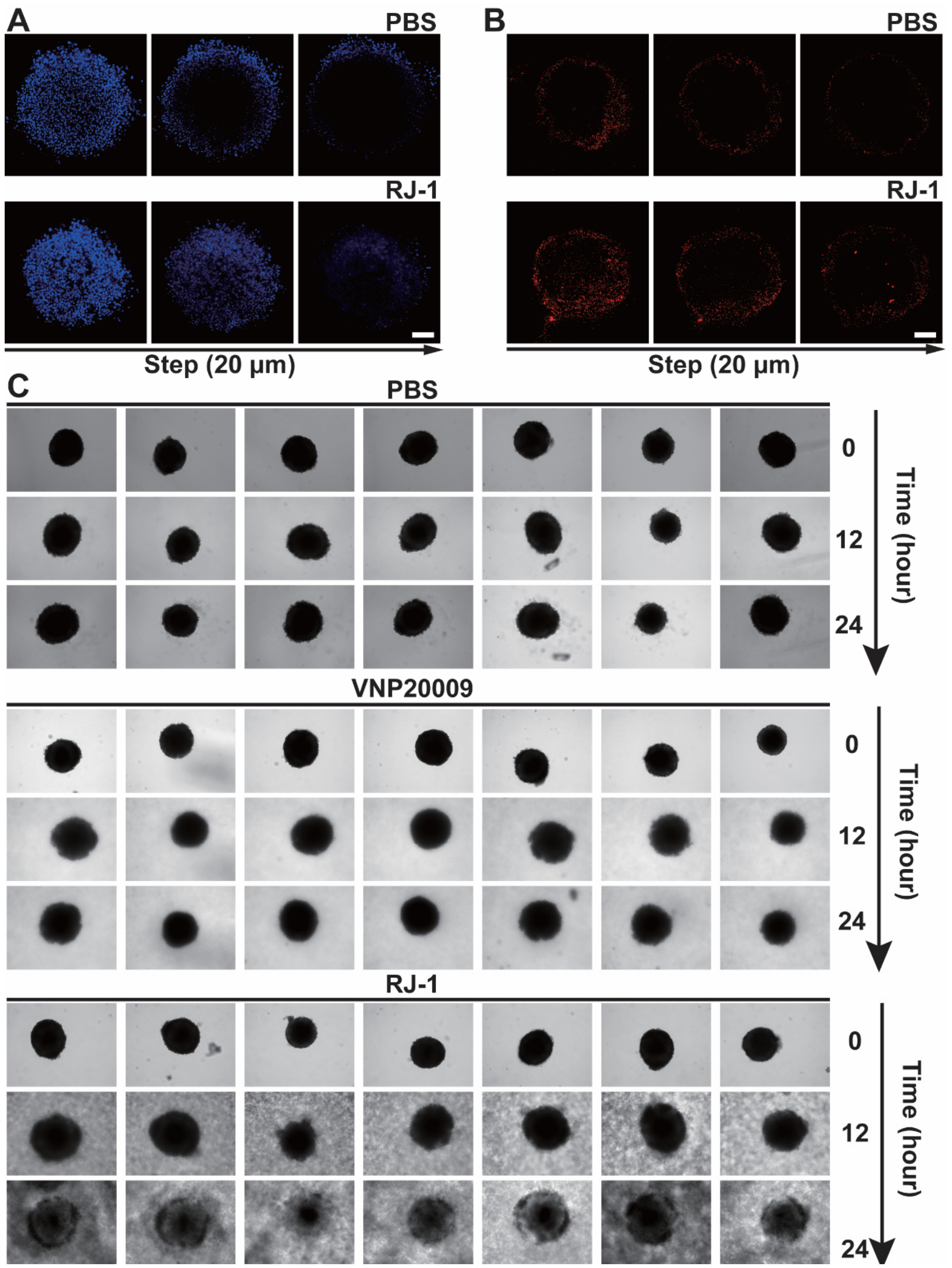


**Fig. S3 RJ-1 mediated drug penetration and oncolysis in vitro.**

(A) Penetration of Hoechst 33342 into tumor spheroids after co-culture with RJ-1 or equivalent volume of PBS visualized by confocal microscopy from bottom to top with a step size of 20 μm (Scale bar: 100 μm). (B) Penetration of liposomes into tumor spheroids after co-culture with RJ-1 or equivalent volume of PBS compared from bottom to top with a step size of 20 μm (Scale bar: 100 μm). (C) Morphological changes of tumor spheroids captured after co-culture with RJ-1, VNP20009 or equivalent volume of PBS for different time periods (Scale bar: 100 μm).

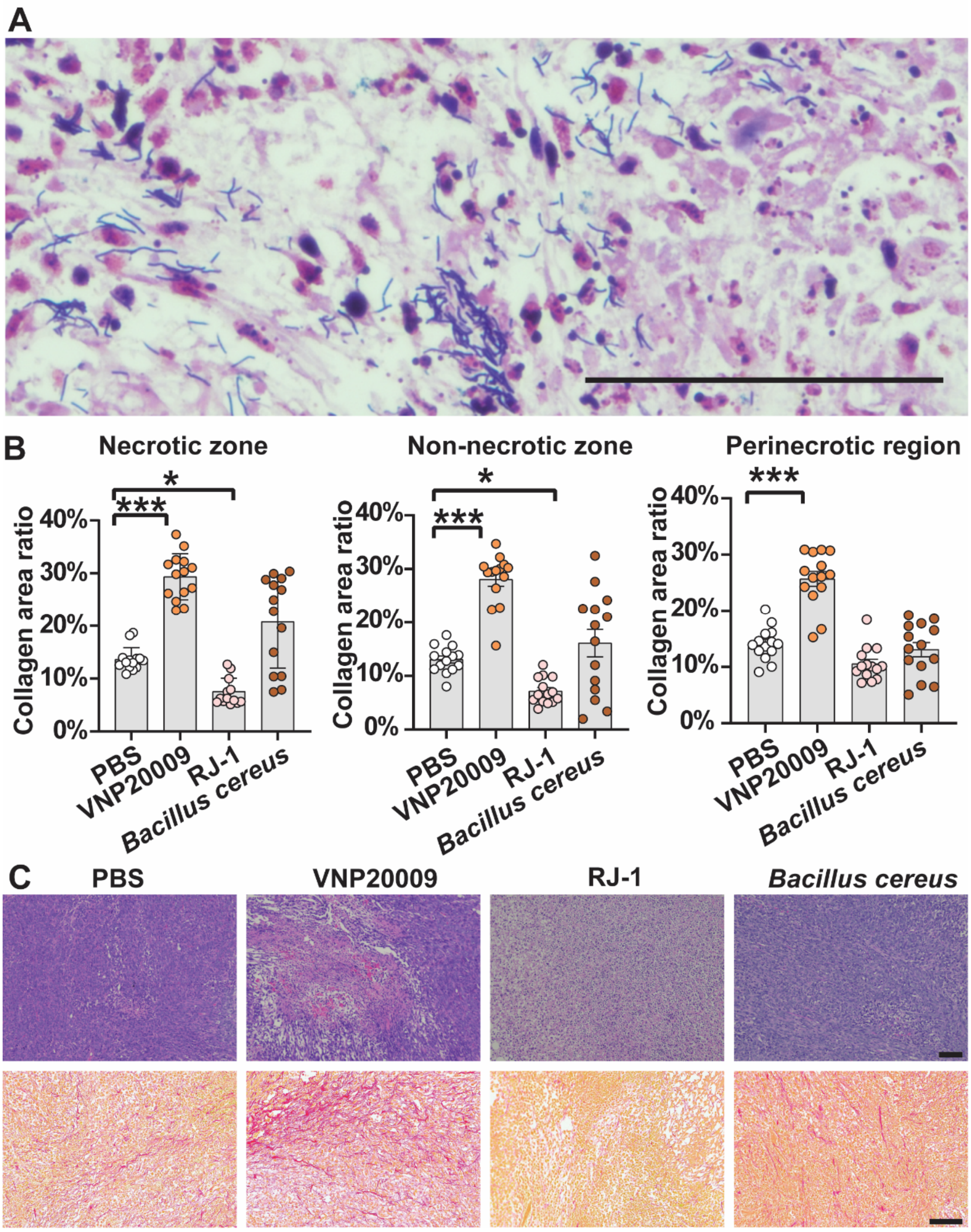


**Fig. S4 RJ-1 survival in TNBC and collagen degradation after intratumoral administration.**

(A) Gram staining of TNBC slices 10 days after treating mice with RJ-1 via intratumoral injection (Scale bar: 100 μm). (B) Proportion of picrosirius red staining area in different regions of tumor slices obtained 10 days after intratumoral injection ($n$ = 14) (Scale bar: 100 μm). (C) H&E staining (top) and picrosirius red staining (bottom) of tumor sections after intratumoral injection of different bacteria or equivalent volume of PBS (Scale bar: 100 μm). For statistical analysis of picrosirius red staining area, one-way ANOVA was used and Tukey test was performed for $p$ value generation. *, $p < 0.05$; ***, $p < 0.001$. Error bar refer to mean value ± SEM.

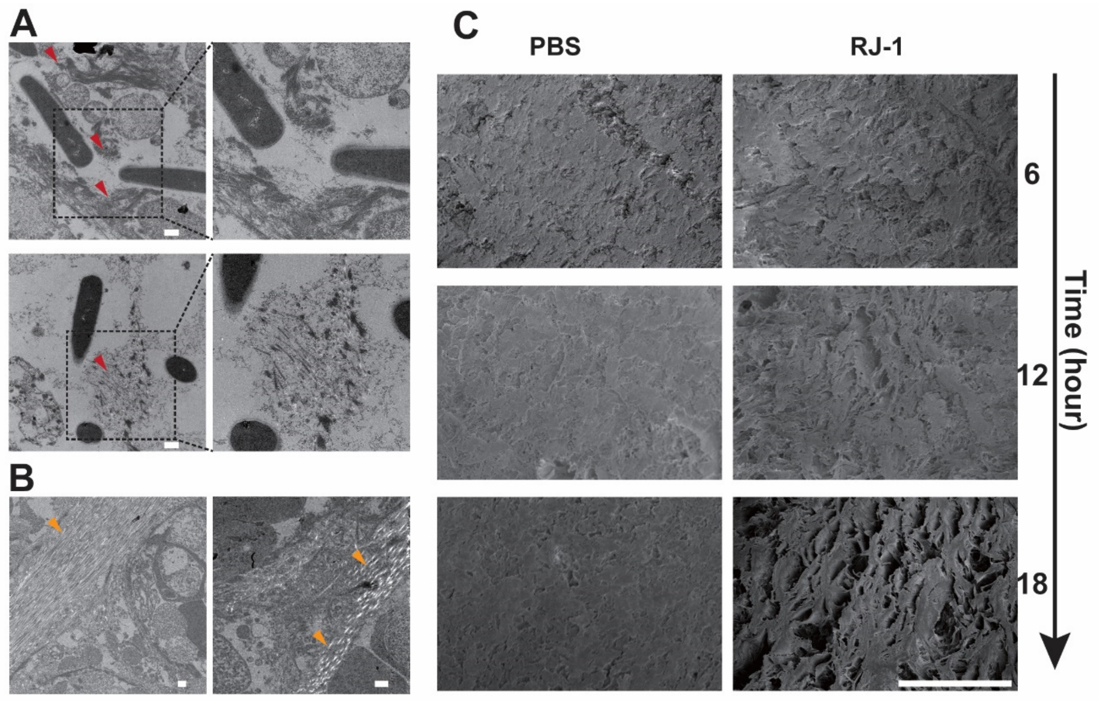


**Fig. S5 RJ-1 mediated disruption of collagen fibers in TNBC.**
(A) Disrupted collagen fibers around RJ-1 vegetative cells. Red arrows refer to digested collagen fibers (Scale bar: 500 nm). (B) Intact collagen fibers in non-necrotic area of the tumor. Yellow arrows refer to intact collagen fibers (Scale bar: 500 nm). (C) Scanning electron microscopy images of collagen mesh in cross sections (Scale bar: 50 μm).

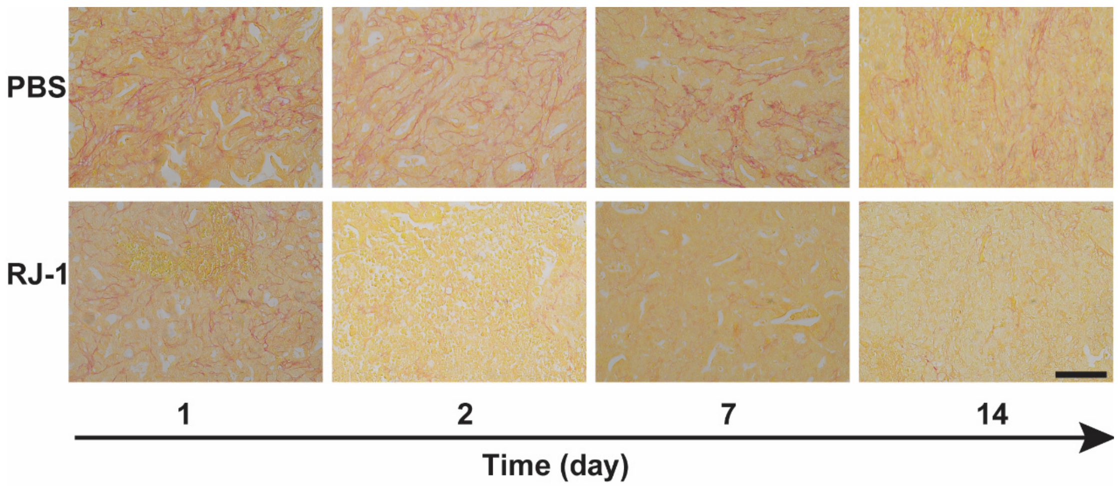


**Fig. S6 RJ-1 mediated degradation of collagen in PDAC.**
Picrosirius red staining was performed in PDAC tumor sections from mice received RJ-1 spore or equivalent volume PBS at different time points (Scale bar: 100 μm).

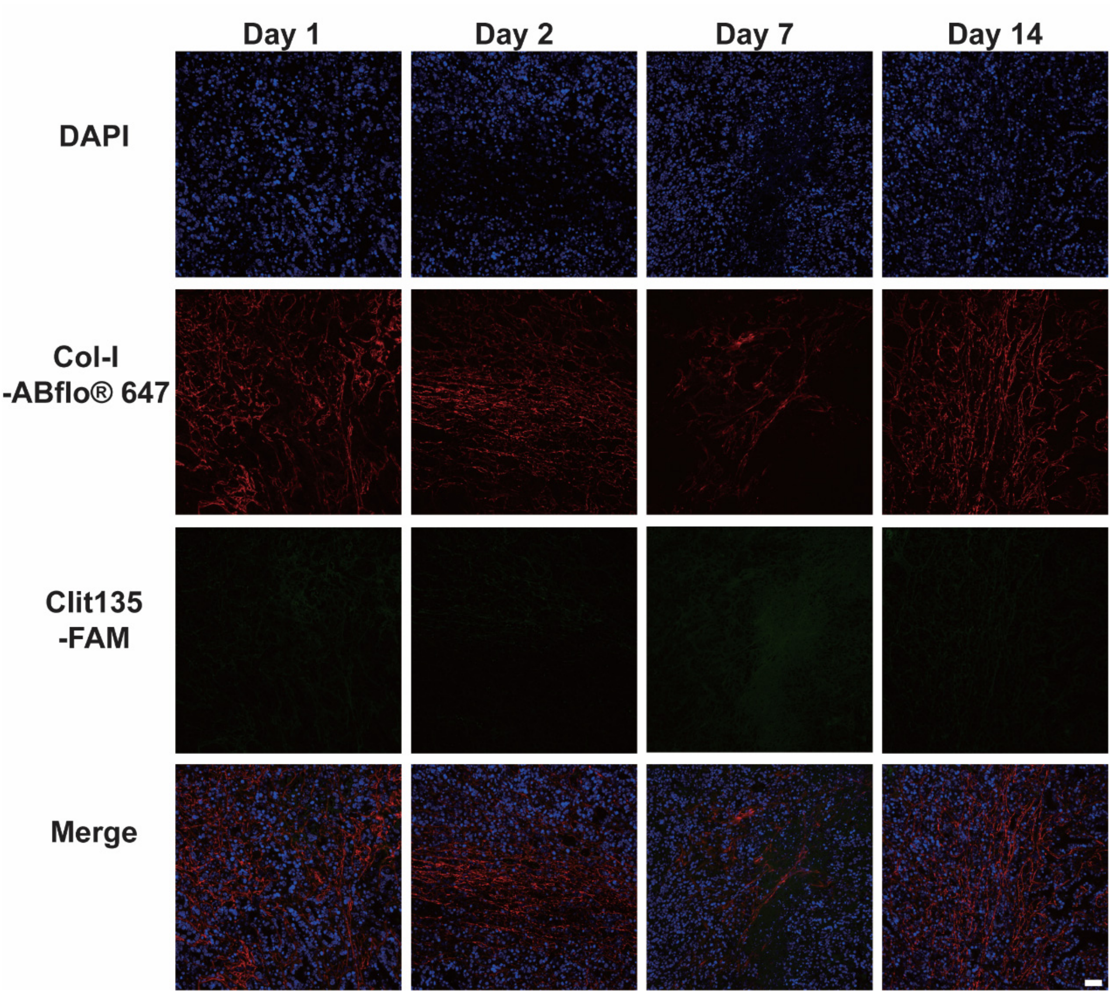


**Fig. S7 Co-localization analysis of Col-I and RJ-1.**

The co-localization of Col-I and RJ-1 was analyzed by immunofluorescence staining of Col-I (red signal) and fluorescence in situ hybridization of RJ-1 (green signal) in PDAC after PBS administration at the indicated time points (Scale bar: 50 μm).

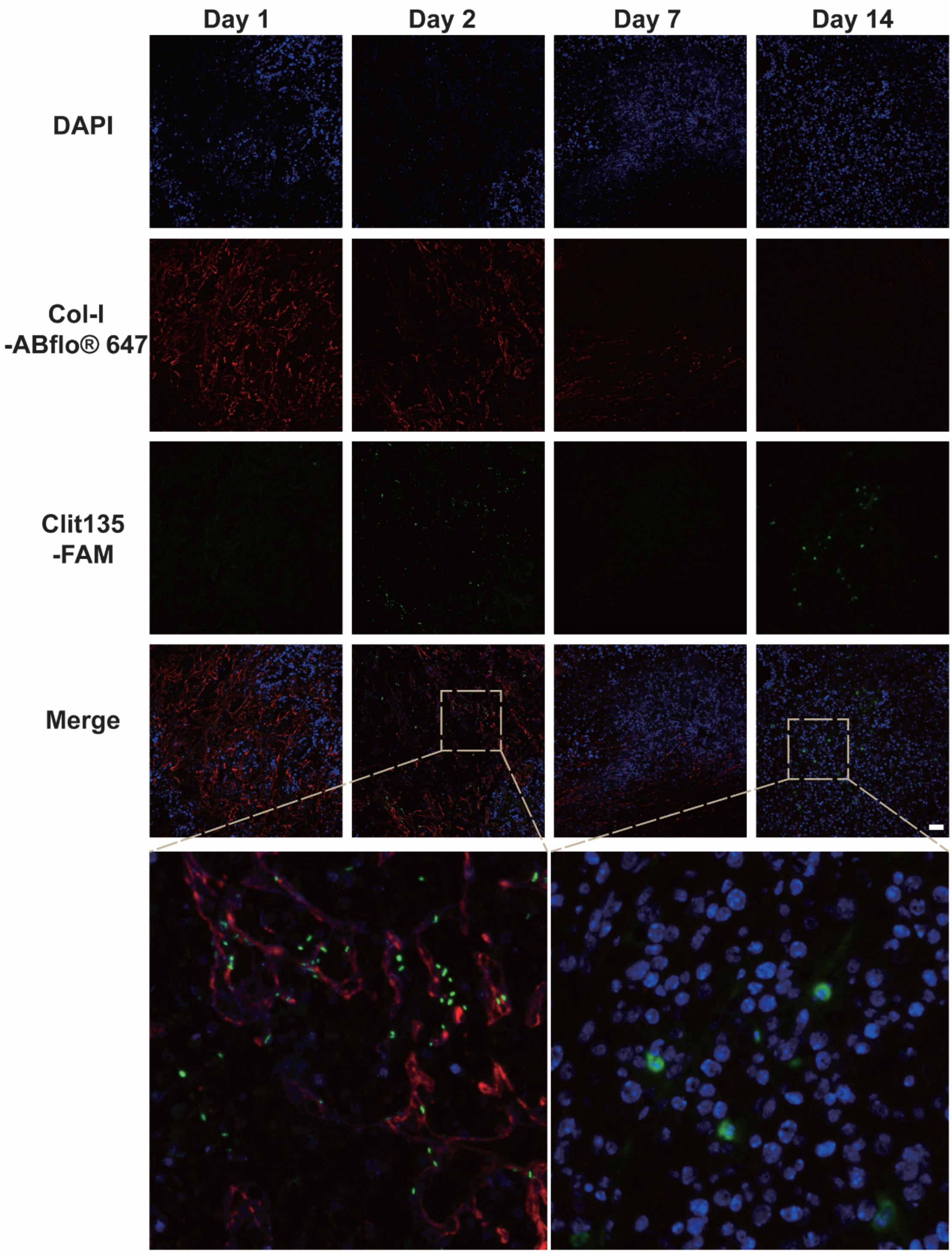


**Fig. S8 Co-localization analysis of Col-I and RJ-1.**

The co-localization of Col-I and RJ-1 was analyzed by immunofluorescence staining of Col-I (red signal) and fluorescence in situ hybridization of RJ-1 (green signal) in PDAC after RJ-1 spores administration at the indicated time points (Scale bar: 50 μm).

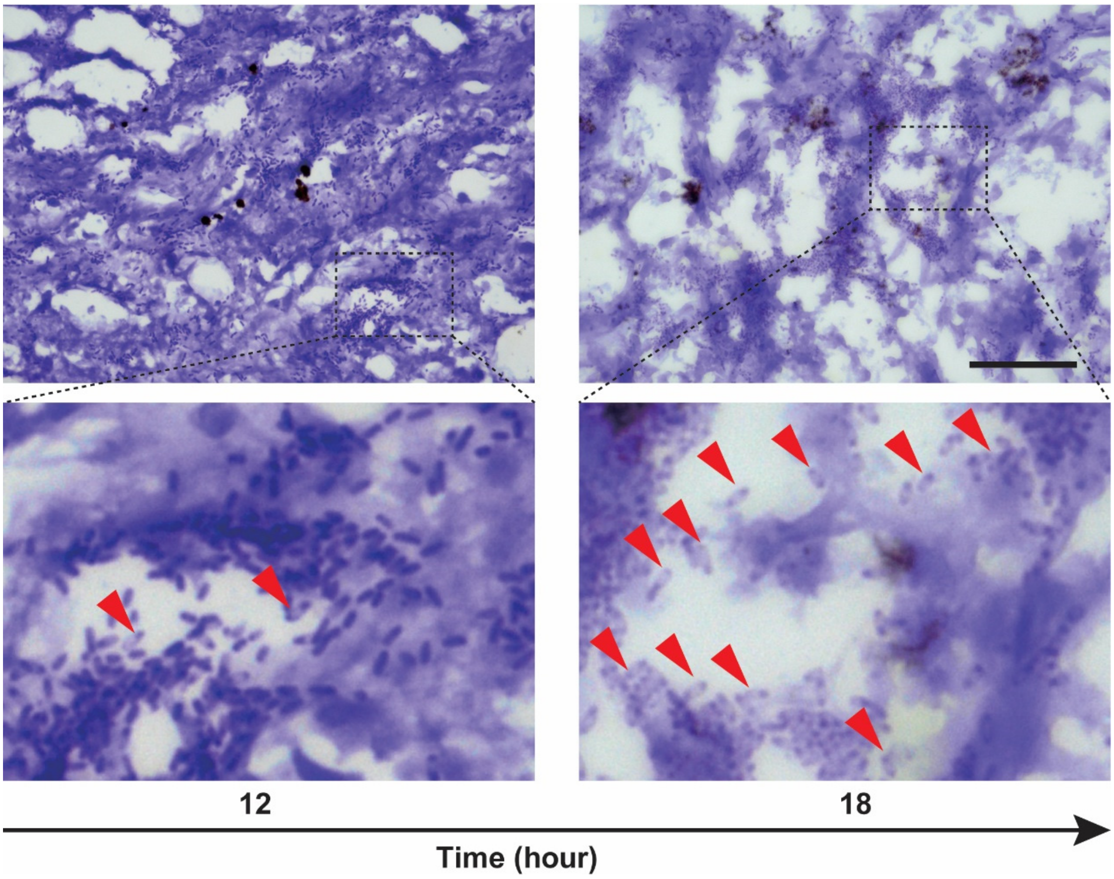


**Fig. S9 Spore formation of RJ-1 in TNBC.**
The re-entry into sporulation process of RJ-1 was observed in TNBC after intravenous injection of $4 \times 10^8$ spores at 18 hours post-injection by Gram staining (Scale bar: 100 μm). Red arrows refer to spores.

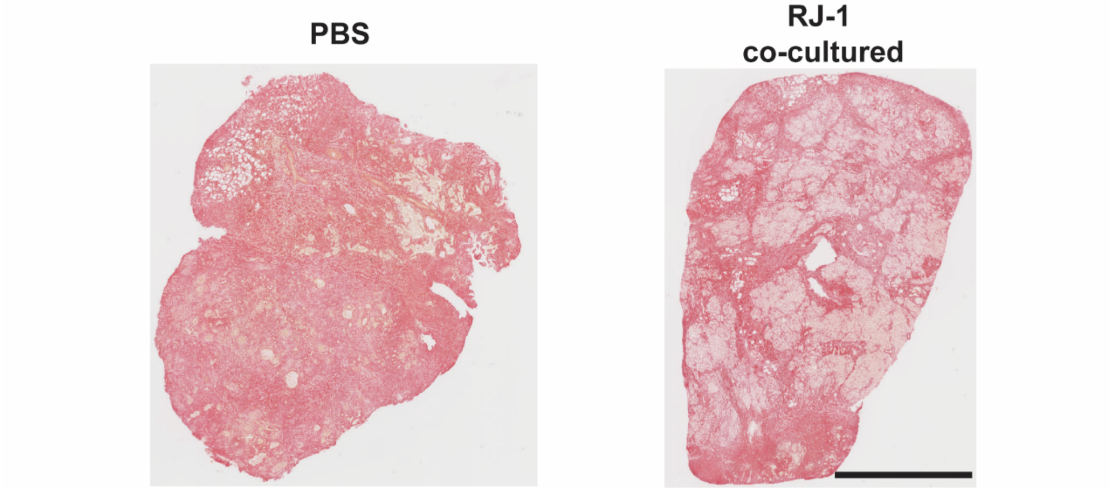


**Fig. S10 RJ-1 mediated collagen degradation in human PDAC tissue.**
Picrosirius red staining was performed in tumor sections from human PDAC that was co-cultured with RJ-1 or equivalent volume PBS for 24 hours (Scale bar: 3 mm).

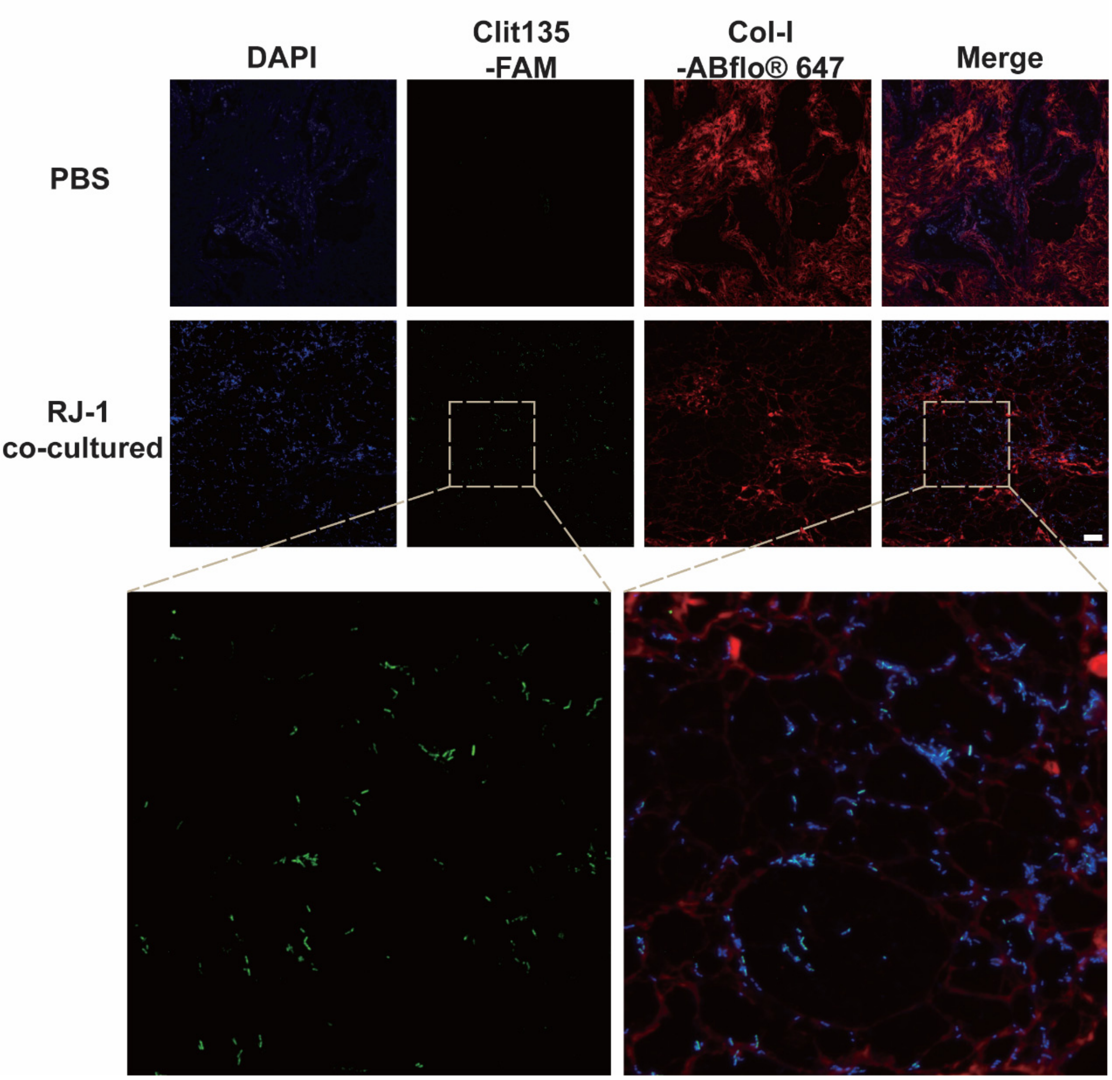


**Fig. S11 RJ-1 germination and Col-I degradation in human PDAC tissue.**

The localization of Col-I and RJ-1 vegetative cells in human PDAC sections were analyzed by immunofluorescence combined with fluorescence in situ hybridization (Scale bar: 50 μm). The rod-shaped structures labeled with Clit-135 represent the vegetative cells, while those not labeled with Clit-135 but labeled with Hoechst 33342 indicate the spores.

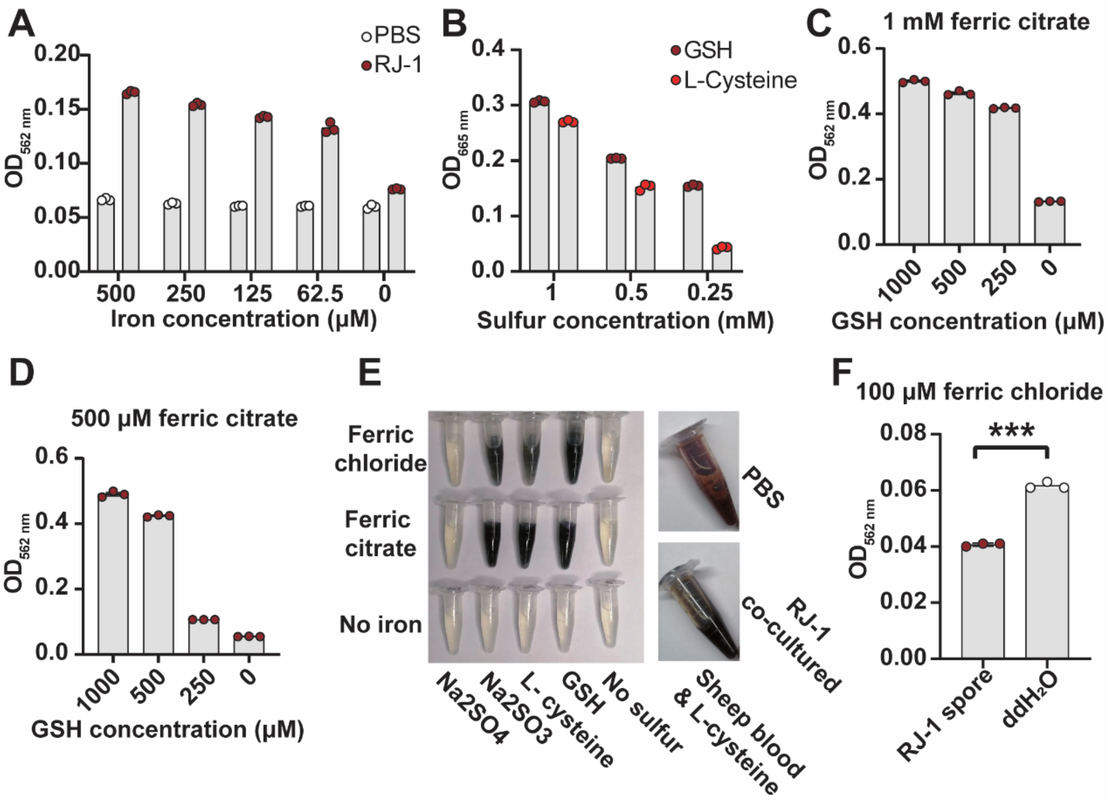


**Fig. S12 In vitro adsorption and mineralization of ferric ion in RJ-1.**

(A) Ferric ion reducing ability of RJ-1 in growth medium. The absorbance was measured at $OD_{562\ nm}$ to quantify the extent of ferrous ions. ($n$ = 3). (B) RJ-1 using different sulfur sources to produce sulfide ions (measured absorbance at $OD_{665\ nm}$). GSH refers to glutathione ($n$ = 3). (C) Sulfur source dose-dependent ferrous sulfide production by RJ-1 at 1 mM iron source concentration. Ferrous sulfide contents were determined by absorbance at $OD_{562\ nm}$ to quantify ferrous ions following acid treatment ($n$ = 3). (D) Sulfur source ferrous sulfide production by RJ-1 at 500 μM iron source concentration. Ferrous sulfide contents were determined by absorbance at $OD_{562\ nm}$ to quantify ferrous ions following acid treatment ($n$ = 3). (E) RJ-1 producing black ferrous sulfide particles in presence of different sulfur and iron sources including defibrinated sheep blood. GSH refers to glutathione. (F) Adsorption of ferric ions by RJ-1 spores or equivalent volume of distilled deionized water ($ddH_2O$). The difference of ferric ion concentration was determined by measuring absorbance at $OD_{562\ nm}$ after being reduced by hydroxylamine hydrochloride ($n$ = 3). For statistical analysis of ferric ions adsorption, we used unpaired student's $t$ test to generate $p$ value. ***, $p < 0.001$. Error bar refer to mean value ± SEM.

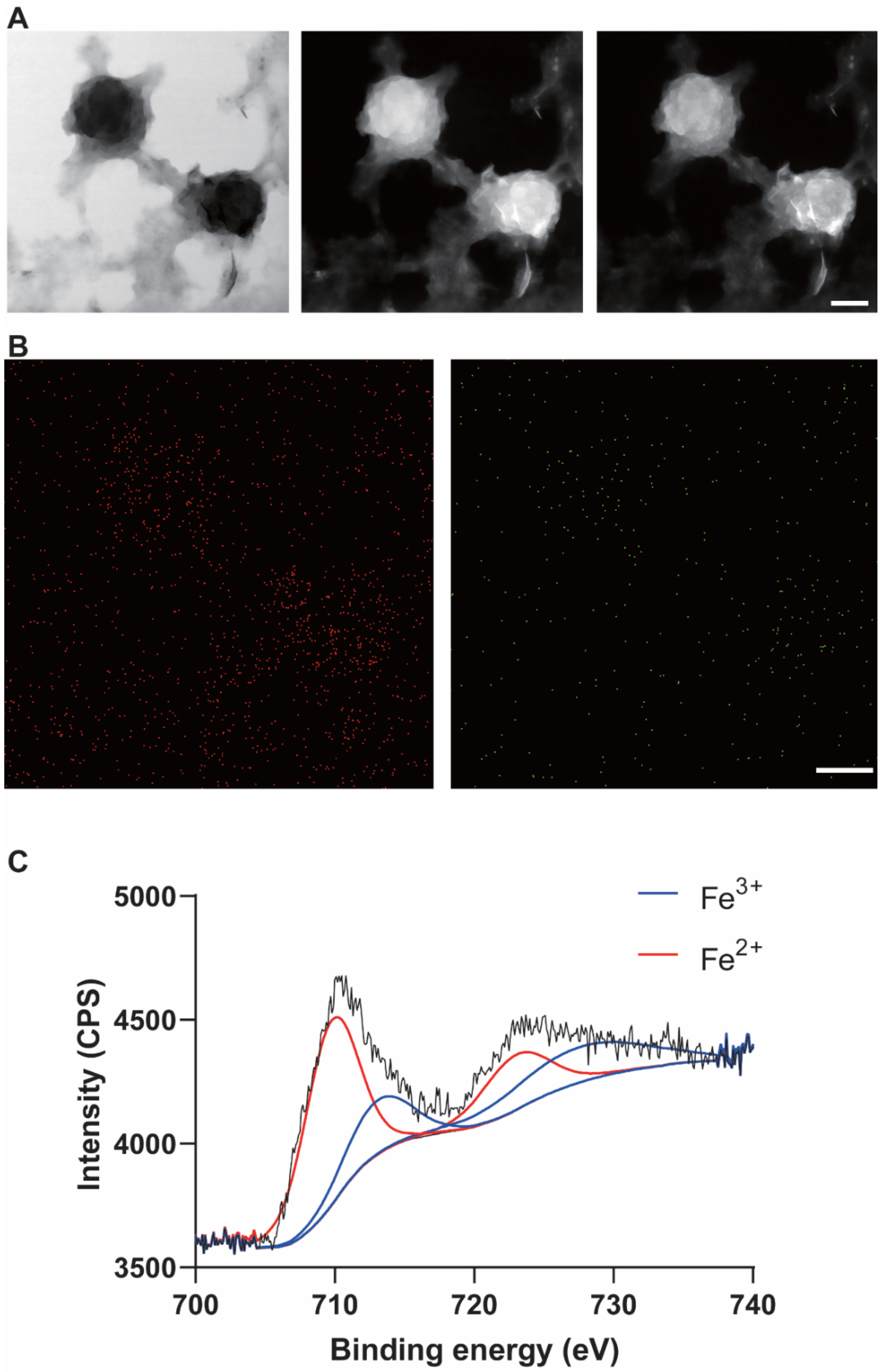


**Fig. S13 Morphology and elemental analysis of INPs.**
(A) Morphology of INPs in bright-field, dark-field, and HAADF modes, along with (B) iron (left) and sulfur (right) distribution displayed in the EDS maps in this field of view (Scale bar: 200 nm). (C) INPs produced by RJ-1 during the cultivation process were collected, dried, and used for X-ray photoelectron spectroscopy analysis of iron.

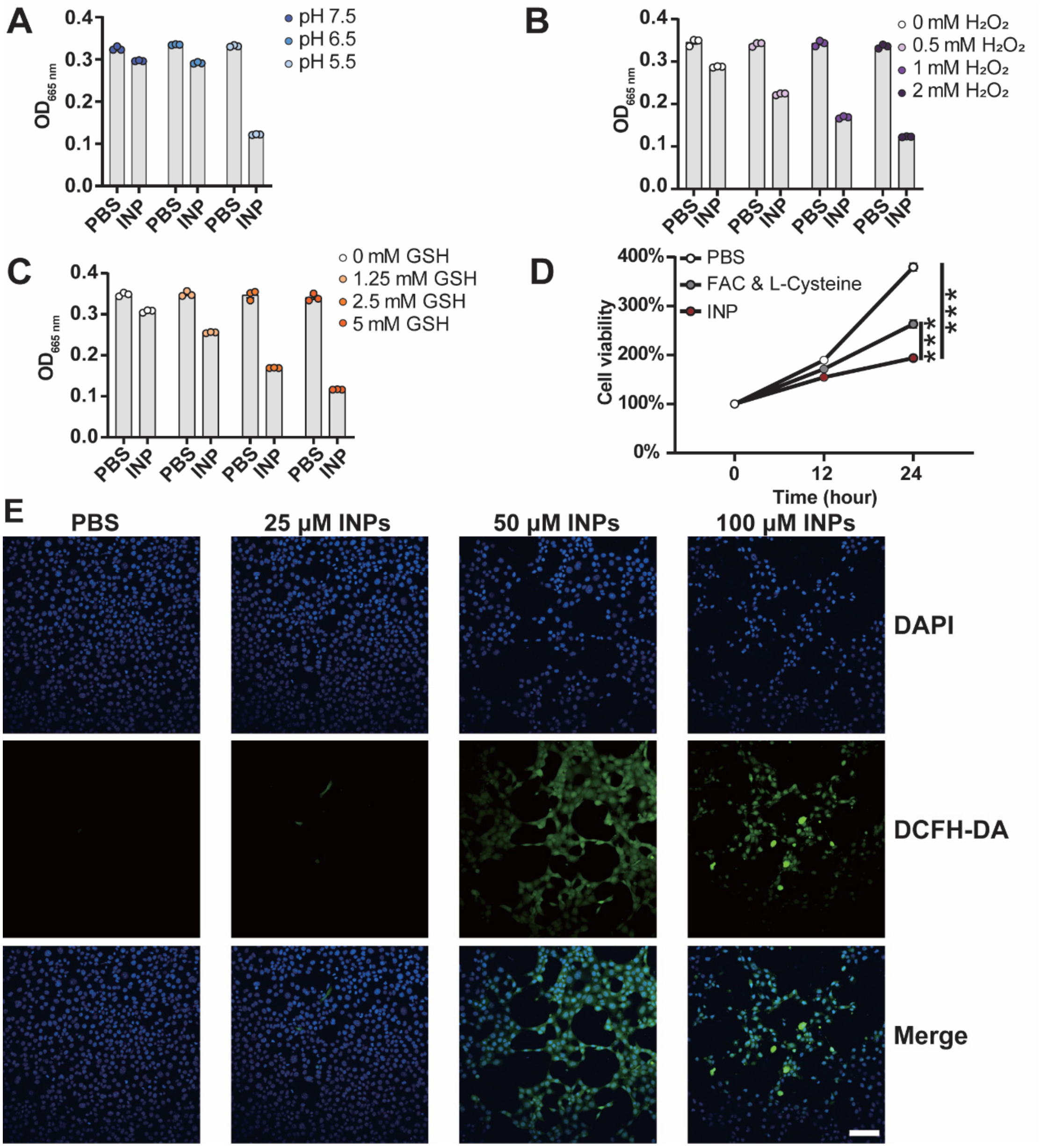


**Fig. S14 INPs induced Fenton reaction in vitro.**

(A) Methylene blue degradation assay verified ROS generation by INPs addition under different pH conditions ($n$ = 3) through the absorbance measurement at $OD_{665\ nm}$. (B) Hydrogen peroxide concentration-dependent production of ROS ($n$ = 3). The absorbance was measured at $OD_{665\ nm}$. (C) GSH concentration-dependent production of ROS ($n$ = 3). The absorbance was measured at $OD_{665\ nm}$. (D) In vitro cytotoxic effects of INPs on 4T1 cells compared to equivalent FAC with L-cysteine or PBS ($n$ = 6). The cells were continuously monitored using Incucyte S3 for a period of 24 hours, and their viabilities were assessed based on the measurement of the total cell area. (E) INPs induced ROS production in a dose-dependent manner on 4T1 cells after 12 hours. DCFH-DA refers to 2',7'-Dichlorodihydrofluorescein diacetate (Scale bar: 100 μm). For statistical analysis cytotoxic effect of INPs, one-way ANOVA was used and Tukey test was performed for $p$ value generation. ***, $p < 0.001$. Error bar refer to mean value ± SEM.

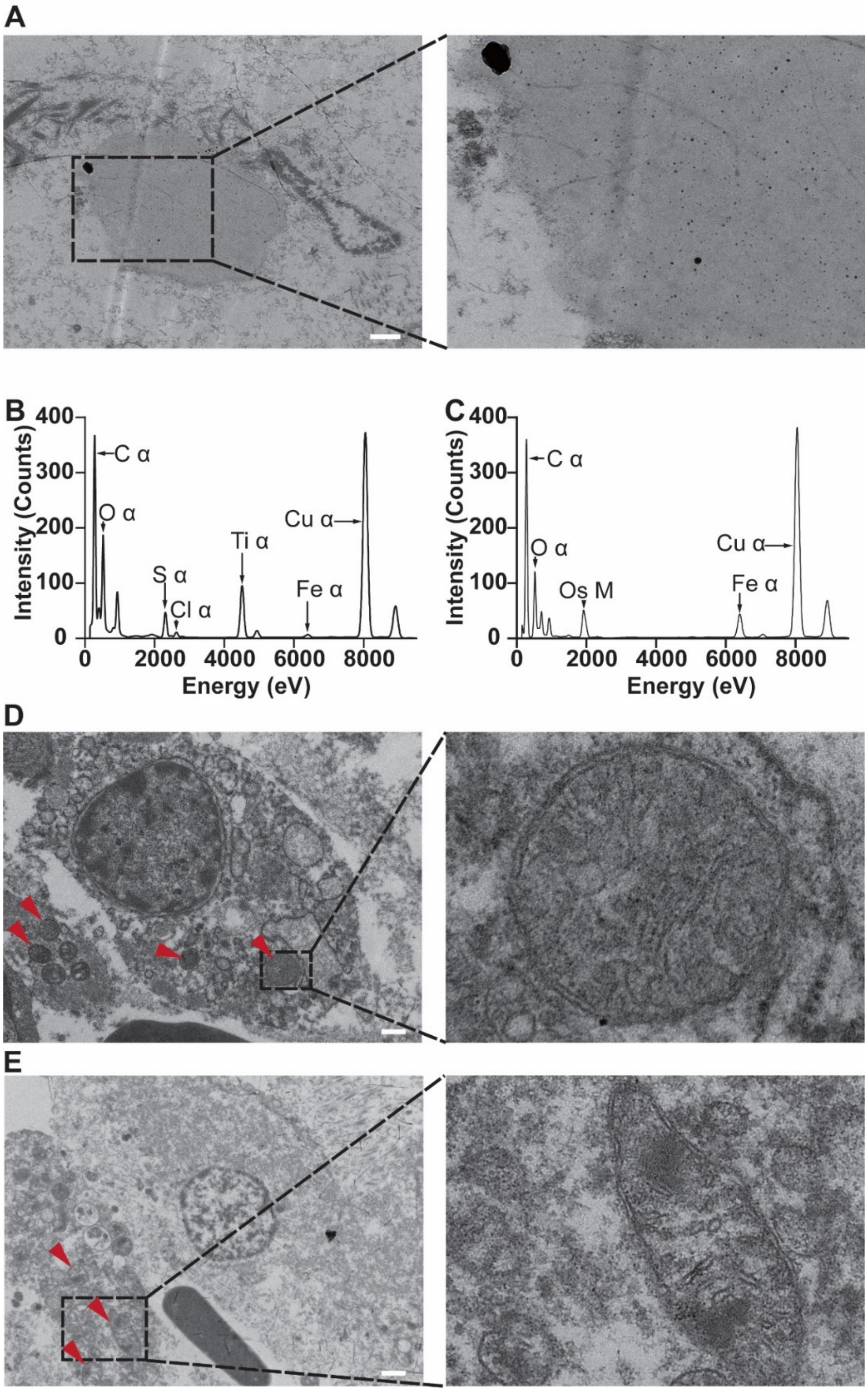


**Fig. S15 EDS analysis of INPs within ultra-thin TNBC sections and ferroptosis morphology of the cells.**

(A) High-electron-density particles released via autolysis (Scale bar: 500 nm). (B) Iron and sulfur signals in INPs, referring to particles shown in Fig. 4G. (C) Iron signal only, referring to particles indicated in Fig. 4H. (D) Red arrows indicate shrunken mitochondria with increased membrane density (Scale bar: 500 nm). (E) Red arrows indicate mitochondria experiencing disruption and loss of cristae (Scale bar: 500 nm).

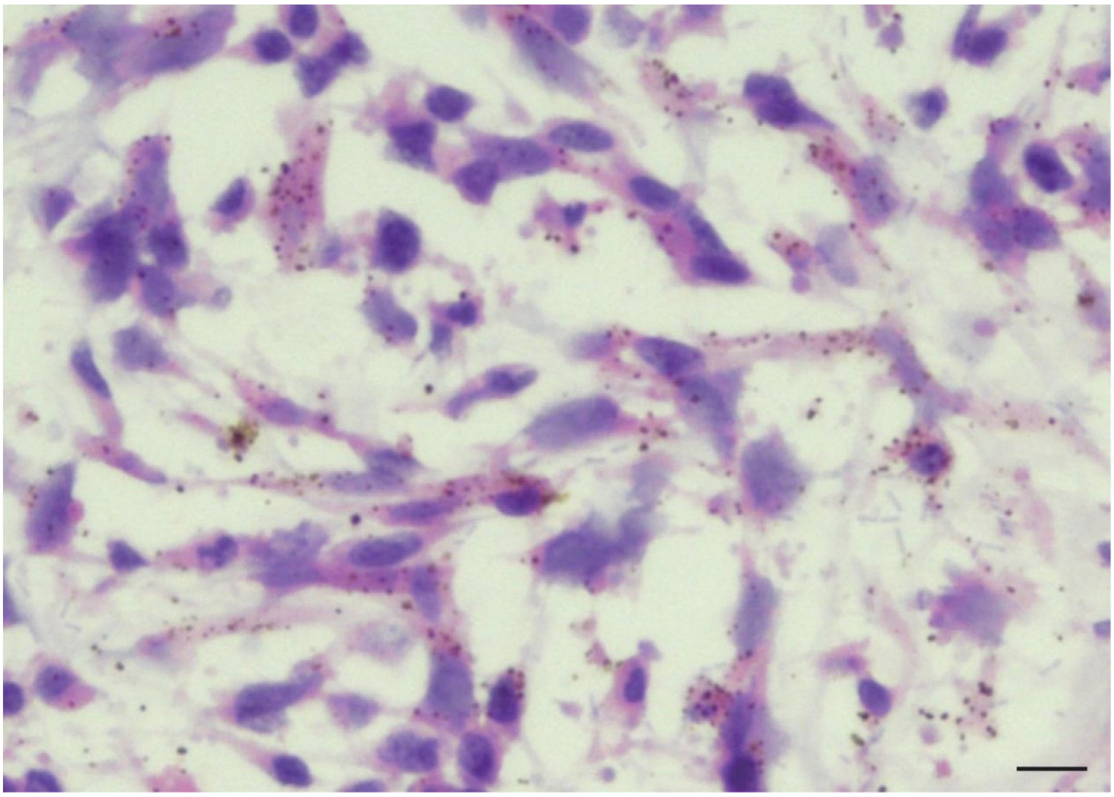

**Fig. S16 Distribution of INPs in viable region of TNBC.**

Distribution of INPs (represented in dark dots) was analyzed by H&E staining 18 hours after intravenous injection with $4 \times 10^8$ RJ-1 spores in TNBC-bearing mice (Scale bar: 10 μm).

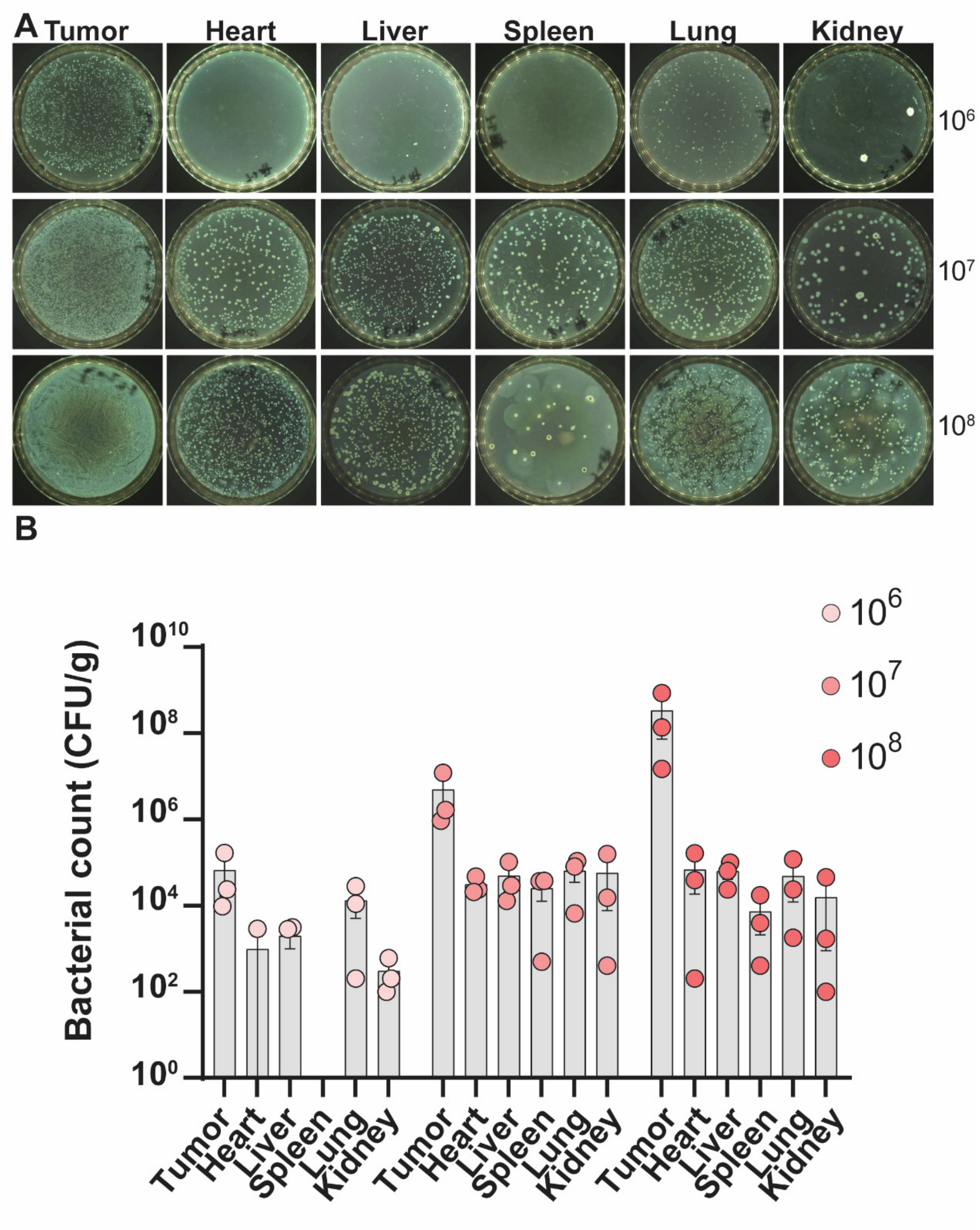


**Fig. S17 Tumor targeting ability of RJ-1.**

(A) Typical images and (B) statistical analysis of RJ-1 bacterial counts in different organs of TNBC-bearing mice 10 days after intravenous injection of different doses of spores. CFU/g refers to colony forming unit per gram tissue ($n$ = 3).

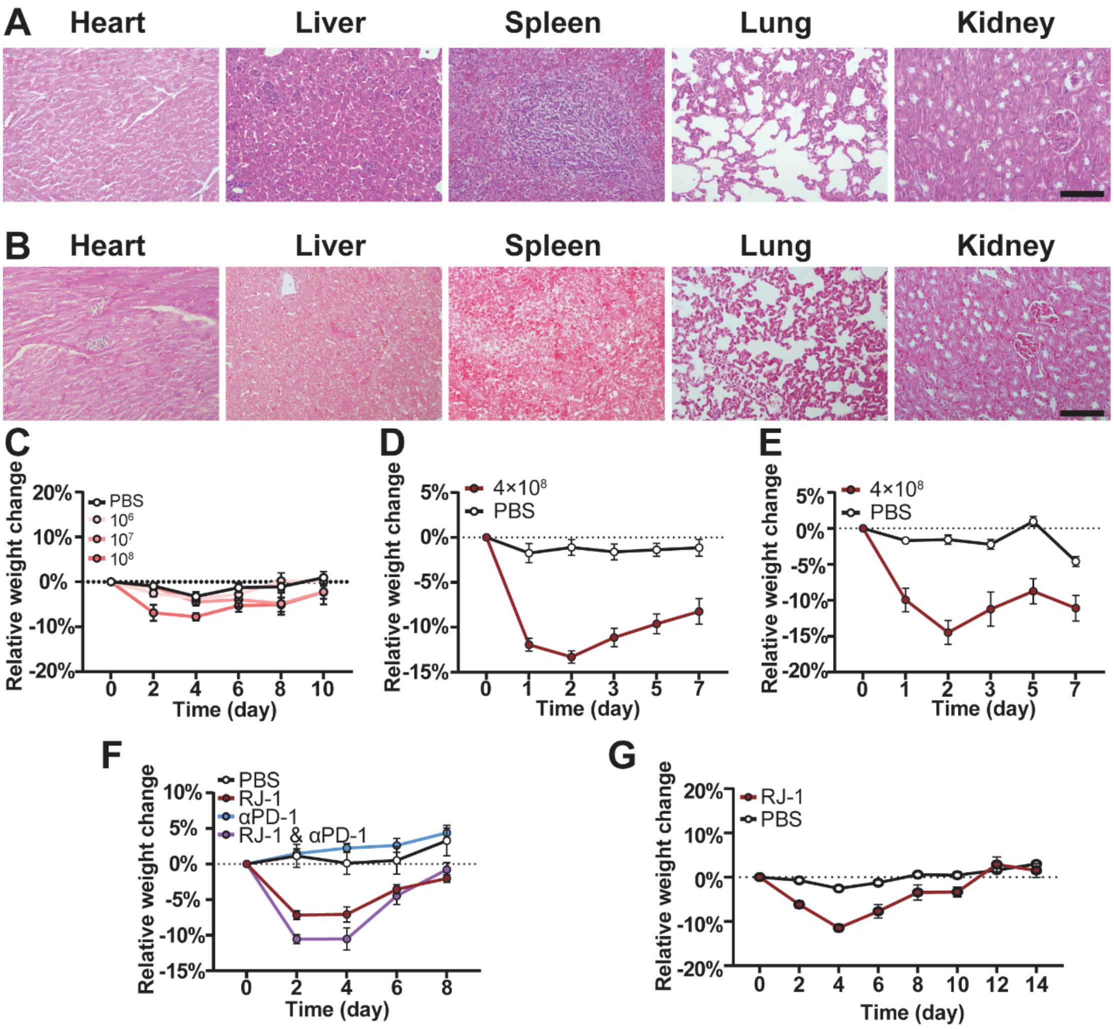


**Fig. S18 In vivo safety of RJ-1.**

(A) H&E staining and (B) Gram staining of tissue sections from different organs of TNBC-bearing mice 10 days after intravenous injection of $10^8$ RJ-1 spores (Scale bar: 100 μm). (C) Body weight changes of mice bearing orthotopic TNBC after intravenous injection of different doses of spores ($n$ = 4). (D) Body weight changes of mice bearing orthotopic TNBC with an average size of ~550 $mm^3$ after intravenous injection of 4 × $10^8$ spores or PBS ($n$ = 7). (E) Body weight changes of mice bearing giant orthotopic TNBC with an average size of ~1000 $mm^3$ after intravenous injection of 4 × $10^8$ spores or PBS ($n$ = 7). (F) Body weight changes of mice bearing orthotopic TNBC after treatment with the combination of αPD-1 and RJ-1 (4 × $10^8$ spores) or equivalent volume of PBS ($n$ = 7-10). (G) Body weight changes of mice bearing orthotopic PDAC after receiving RJ-1 spores ($10^9$) or PBS ($n$ = 6). Error bar referred to mean value ± SEM.
